# Glutathione accelerates the cell cycle and cellular reprogramming in plant regeneration

**DOI:** 10.1101/2023.11.28.569014

**Authors:** Laura R Lee, Bruno Guillotin, Ramin Rahni, Chanel Hutchison, Bénédicte Desvoyes, Crisanto Gutierrez, Kenneth D Birnbaum

## Abstract

The plasticity of plant cells underlies their wide capacity to regenerate, with increasing evidence in plants and animals implicating cell cycle dynamics in cellular reprogramming. To investigate the cell cycle during cellular reprogramming, we developed a comprehensive set of cell cycle phase markers in the Arabidopsis root. Using single-cell RNA-seq profiles and live imaging during regeneration, we found that a subset of cells near an ablation injury dramatically increases division rate by truncating G1. Cells in G1 undergo a transient nuclear peak of glutathione (GSH) prior to coordinated entry into S phase followed by rapid divisions and cellular reprogramming. A symplastic block of the ground tissue impairs regeneration, which is rescued by exogenous GSH. We propose a model in which GSH from the outer tissues is released upon injury licensing an exit from G1 near the wound to induce rapid cell division and reprogramming.

## Introduction

Plants have a remarkable capacity to regenerate, where even a single somatic cell can give rise to an entire organism^1^. The *Arabidopsis* root apical meristem (RAM) provides a model for regeneration in plants because the organ can regenerate from differentiated cells after removal of the stem niche without exogenous hormones^2^. This process requires the coordination of both cell division and cell identity changes among the cells that will give rise to the new, regenerated tissue. How cell division properties are coordinated with cell fate change in regeneration remains an open question.

Prior work in plants has demonstrated links between cell cycle regulation and cell fate specification. In the *Arabidopsis* sepal, giant cells are specified when *MERISTEM LAYER 1* (*ML1*) expression exceeds a threshold level during G2/M phase of the cell cycle^3^. Recent work has shown that high protein levels of the cell fate regulators SHORT ROOT (SHR) and SCARECROW (SCR) at a specific phase of the cell cycle determine the polarity of a formative division in the root^4^. In the stomatal lineage, asymmetric and symmetric cell divisions are mediated by the expression of a series of master regulator basic helix-loop-helix (bHLH) transcription factors that concomitantly regulate cell identity (reviewed in^5^).

After specification, cell cycle length frequently changes as cells undergo differentiation. For example, in the root meristem, cell division rates speed up along the maturation gradient as they move away from the stem cell niche during transit amplification^6^, largely due to a shortening of G1 duration as a cell is displaced shootward away from the quiescent center (QC) ^7^. Alternatively, in the stomatal lineage, G1 duration increases and cell cycles lengthen as cells commit to terminal differentiation^8^. These observations suggest that even the trends of cell cycle length dynamics during differentiation can differ from tissue to tissue.

Different specialized plant cells can also show differences in their core cell cycle machinery. While many core cell cycle regulators are conserved between plants and animals^9,10^, the expansion of core cell cycle gene families in plants, such as the D type cyclins, has allowed for cellular specialization^11^. For instance, *CYCLIN D-6* (*CYCD6*) is specifically expressed downstream of the SHR-SCR module and mediates the switch in division orientation that leads to the formation of a new cell type^12^. *CYCLIN D-7* (*CYCD7*) expression is restricted to the guard mother cell in the stomatal lineage and regulates a switch from asymmetric to symmetric divisions^13^. This process is also regulated by another specialized cyclin, *CYCLIN D-5* (*CYCD5*)^14^.

These examples suggest that cell type-specific regulation of the cell cycle in plants could be a more general phenomenon, but one challenge facing the field has been studying the cell cycle in a way that maintains the developmental context of cells. Early transcriptional studies of the cell cycle in *Arabidopsis* employed synchronization of cultured cells^15,16^, which provided valuable insight, but such bulk, *in vitro* analysis could not provide cell type-specific information.

Single-cell RNA-seq (scRNA-seq) studies provide an opportunity to analyze cell-type specific properties of the cell cycle, but, outside of G2/M, the ability to map cells to a cell cycle phase is limited. For example, the field lacks a reliable transcriptional marker for the G1 phase of the cell cycle, although CDT1a is a well-supported translational marker^17,18^. Overall, we have an incomplete view of the extent of cell cycle variation among cells and the transcriptional signatures that distinguish the different ways the cell cycle varies in specific cell types. An extensive set of markers for each cycle phase would add insight into the analysis of cell cycle attributes in the burgeoning collection scRNA-seq studies in plants.

In the context of regeneration, cell division is required for complete repair of injured tissues^2^. There is considerable evidence in animals that events during the G1 phase of the cell cycle are critical for cell fate establishment^19–21^, and short G1 phases are a known feature of totipotent animal stem cells^22^. In plants, division rates can vary dramatically in different contexts. For example, cell divisions in the transit amplifying zone of the root in one study were shown to have a median duration of 21.5 hr^6^. However, other studies showed that cells divide approximately every 7 hr in Arabidopsis embryos up until the 16 cell stage^23^, and in lateral root primordia cell doubling time has been estimated to be as short as 3.7 hr^24^. It is not known in plants if rapid divisions facilitate organ formation or cell fate specification in any of these contexts, including regeneration.

Both plants and animals can vary division rates by controlling the passage through G1 and G2 checkpoints, which are often regulated by metabolites^9,25^. For example, it was recently shown that tricarboxylic acid cycle metabolites may regulate root growth and development^26^. In addition, reactive oxygen species (ROS) have a role in controlling division rates along the root axis^27^. The tripeptide glutathione (GSH) – the primary antioxidant in the cell^28^ – is enriched in the nucleus in division-competent cells in both plants and animals with the hypothesized function of protecting newly synthesized DNA from ROS-induced damage (reviewed in^29^).

GSH availability^30,31^ and ROS patterning^32^ in plants have previously been linked to root growth and cell cycle regulation^30,32,33^. Prior studies have shown that GSH may be necessary for plant cells to pass the G1 to S transition^30^ and nuclear ROS levels change cyclically in cell cycle-synchronized root tips^34^. Finally, evidence from Arabidopsis tissue culture suggests that GSH is transported into the nucleus in a cell cycle-dependent manner with consequences for the redox state of both the nucleus and the cytoplasm^33^. Injury, such as RAM excision, results in accumulation of ROS that presumably need to be neutralized by antioxidants^35^. While both GSH availability and cell cycle regulation are linked to cellular reprogramming following injury, how these factors are coordinated during regeneration, if at all, remains unknown.

Here we generate transcriptomic profiles of the cell cycle in the RAM while maintaining developmental context. We synchronize cells in the intact RAM^36^ followed by scRNA-seq to generate phase-enriched profiles for the G1, S, and G2/M phases of the cell cycle. We corroborate these transcriptional profiles with phase-specific bulk RNA-seq profiles and *in situ* hybridizations to yield a high-confidence set of cell cycle markers. In conjunction with scRNA-seq profiles, we used the markers to analyze the transcriptional composition of each phase of the cell cycle both broadly and among specific cell types. Collective analysis of these datasets reveals 1) many individual cell types have distinct cell cycle dynamics at the transcriptional level and 2) the G1 phase of the cell cycle is uniquely tuned to respond to redox stress. During regeneration, we used both single-cell analysis and live imaging to show a dramatic shortening of the G1 phase of the cell cycle in cells near the injury. Furthermore, cells with a short G1 phase reprogram to new cell fates more rapidly than neighboring cells that maintain a longer G1. We demonstrate that GSH mediates both the rapid exit from G1 and fast divisions that preferentially lead to cellular reprogramming. Finally, the results showed that the middle and outer cell types appear to be a major source of GSH in the root that facilitates growth and regeneration. Overall, we show that GSH acts as a signal in regeneration where, upon wounding, GSH enters the nucleus, prompting a rapid exit from G1, a fast cell cycle, and cell-fate reprogramming. Our work establishes a new role for GSH as an injury communication molecule that regulates cell cycle duration to mediate organ regeneration.

## Results

### Phase-enriched scRNA-seq libraries reveal a large set of cell cycle-regulated genes

To gain a deeper view of cell cycle dynamics in specific cell types, we sought to characterize cell cycle transcriptomes in intact Arabidopsis roots while maintaining developmental context. To that end, we synchronized cells *in* vivo using hydroxyurea (HU)^36^ followed by scRNA-seq to obtain phase-enriched populations in which cell type-specific information is maintained (Figures 1A and 1B, Figures S1-S2). We then performed a differential expression analysis among the phase-enriched libraries as implemented in Seurat^37^,, asking for only positive markers that passed a p-value threshold of < 0.01 to identify genes specific of each phase-enriched library. To prevent any cell type from contributing an excess of markers to the final set, we selected the same number of cells per cell type in each phase-enriched library (Figure S3A,B). For each marker we calculated the difference between the percentage of cells expressing the given marker in the library in which it was differentially upregulated and the percentage of cells expressing that marker in all other libraries. We ranked each phase marker based on this calculation and selected the top 50 genes from each set to further ensure the selection of high confidence cell cycle markers (Table S1). For example, a gene shown to be upregulated in the G1-enriched library was ranked more highly if it was also expressed in a higher proportion of cells in the same G1-enriched library. This approach identified gold standard cell cycle markers in their appropriate phase (Figure S3C, Table S2).

**Figure 1:**
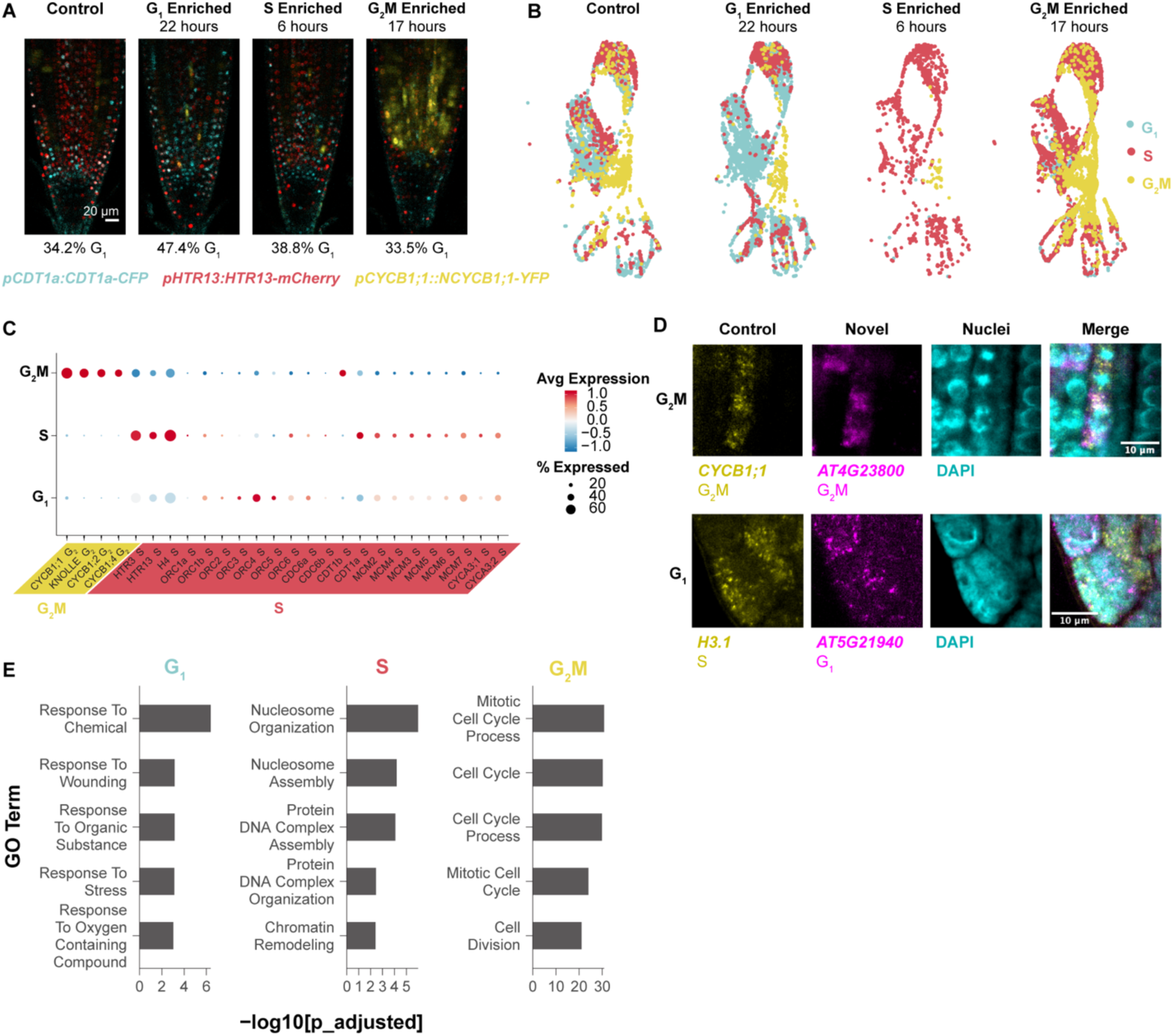
Single-cell phase synchronized cells yield robust transcriptional markers for each phase of the cell cycle. (A) Representative images of phase enrichments achieved with HU synchronization using seedlings expressing the PlaCCI reporter. The G1 phase reporter *pCDT1a:CDT1a-CFP* is shown in cyan, the S phase reporter *pHTR13:HTR13-mCherry* is shown in red, and the G2/M phase reporter *pCYCB1;1:NCYCB1;1-YFP* is shown in yellow. The percentage of cells in G1 (CFP positive cells/All cells), determined from 3D segmentation of z stacks, is shown below each panel. The hours noted above each panel refer to the treatment time each batch of roots was exposed to 10mM HU for. (B) An unsynchronized control and three single cell profiles collected at specific times after synchronization integrated in UMAP with cells color coded by phase determination. Cells from each time point were separated after integration. (C) A dot plot showing expression of gold standard cell-cycle phase markers, showing known G2/M-phase markers (CYCBs) followed by known S-phase markers. (D) *In situ* hybridization of novel G1 and G2/M probes. Known markers are shown in yellow and new markers in magenta. The new G2/M marker is hybridized with a known G2/M marker, showing overlap. The new G1 marker is hybridized with a known S marker, showing spatial anticorrelation. (E) The top five most statistically significantly enriched GO terms among the top 50 phase marker set for each phase. See also Figures S1-S5 and Tables S1-S3.

To corroborate the markers and account for the potential effects of synchronization, we also generated bulk RNA-seq libraries from root cells sorted by ploidy as a proxy for cell cycle phase^38^. The resulting dataset was analyzed by K-means clustering to reveal the expression patterns of highly variable genes among cell cycle phases. We then analyzed the overlap in cell cycle phase markers between the synchronized single scRNA-seq and bulk RNA-seq profiles (Figure S4). These two approaches overwhelmingly assigned genes to the same phase of the cell cycle (Figure S4).

We then used the stringent top-50 marker set to assign our synchronized cells to phases in Seurat^39^ (Figure 1B) and examined the expression of known G2 and S-phase marker genes with functional roles in the cell cycle (Figure 1C). Canonical G2/M markers – cyclin Bs – and the S-phase markers – minichromosome maintenance complex (MCM) – were classified to the appropriate phase. The origin recognition complex (ORC) family, which is required in S-phase to license DNA replication, appeared to be expressed more highly in G1. This is consistent with the observation that ORCs are required in the pre-replication complex prior to MCMs (reviewed in^40^) and supports that this set of phase markers provides a sensitive discrimination between G1 and S phase.

To validate these markers *in vivo*, we visualized transcripts directly using multicolor *in situ* hybridization (Figure 1D, Figures S3D-S3G). This allowed us to simultaneously visualize a known phase marker in the same plant as well as a novel probe. We observed a novel G2/M marker - AT4G23800 - co-staining with a probe for the well-known G2/M marker CYCB1;1. Additionally, both markers were present in cells with mitotic figures, as visualized by DAPI, further confirming the novel marker is expressed in cells in G2/M phase.

To validate a novel G1 probe, we co-stained a new, putative G1 marker - AT5G21940 - with a well-established S phase marker - AT5G10390 (H3.1)^41^, testing for anticorrelation and exclusion from mitotic figures since there are no known G1 transcriptional markers in plants. As predicted, we found the transcripts of these two genes were both anticorrelated and absent from mitotic figures, with occasional overlap (Figure 1D; Figures S3D-S3G). Thus, the marker set provides a highly sensitive tool for cell cycle analysis in single cell studies and, importantly, a method to distinguish cells in the G1 phase, allowing new analysis of the role of G1 in plants.

The analysis showed that markers for G1 and S phases had expression patterns that were enriched in but not strictly exclusive to their respective phase. For example, S-phase markers, while most highly expressed in that phase, often had low levels of expression in G1 and vice versa. While G2/M was transcriptionally distinct, G1 and S had more continuous expression patterns. However, the full set of markers for each phase, including G1, robustly assigned root cells to a specific phase in scRNA-seq datasets.

The large, high-confidence set of markers allowed us to analyze functional categories in each phase, particularly G1, which has not been deeply characterized. First, as expected, gene ontology (GO) enrichment analysis revealed that cell cycle-related terms were enriched in G2/M and S phases (Figure 1E, Table S3). We found that canonical cell cycle markers are lowly expressed so the most robust markers did not include cyclins (Figure S5). Notably, many markers were enriched in, but not necessarily specific to, any given cell cycle phase, showing that, beyond the distinct transcriptome of G2/M, other phases had less discrete transitions at the transcriptional level.

Interestingly, the top 50 G1 markers were enriched for GO terms related to stress responses (Figure 1F) and closer inspection of a larger marker set showed these terms were specifically related to oxidative stress (Table S3). This enrichment of ontology terms in G1 cells was also present in the G1 ploidy sorted dataset, where cells from each phase were selected from the same pool of cells, ruling out a batch effect (Table S3). This suggests a role for oxidative stress management within G1—an intriguing feature that we pursued below in the analysis of the cell cycle in regeneration. Overall, the dataset now provides a robust tool to analyze the cell cycle in single-cell profiles and identifies new genes with potential roles in specific phases of the cell cycle.

### Pseudotime analysis reveals cell cycle variation within and between cell types

We sought to generate a fine-grained analysis of cell type-specific cell cycle patterns in the Arabidopsis root. Using a scRNA-seq profile of all cells in the root meristem^42^, cells were aligned in a pseudotime trajectory with Monocle3^43^ using only the top 150 cell cycle markers and visualizing them in Uniform Manifold Approximation and Projection (UMAP^44^, Figure 2A). The trajectories were anchored in G2/M and proceed to G1 where they split into three separate branches that each continue to S phase. To map cell identities onto cell cycle-annotated single cells, we used cell-identity markers identified in an independent analysis^42^. Despite filtering out cell type-specific markers in the original ordering, different cell types still favored—but were not restricted to—distinct regions of the UMAP space (Figure 2B, Figure S6). This indicated that, using only a core set of shared cell cycle markers, cells still clustered by their *in vivo* identity, suggesting the separate branches for one cell cycle phase represented cell type-specific cell cycles.

**Figure 2:**
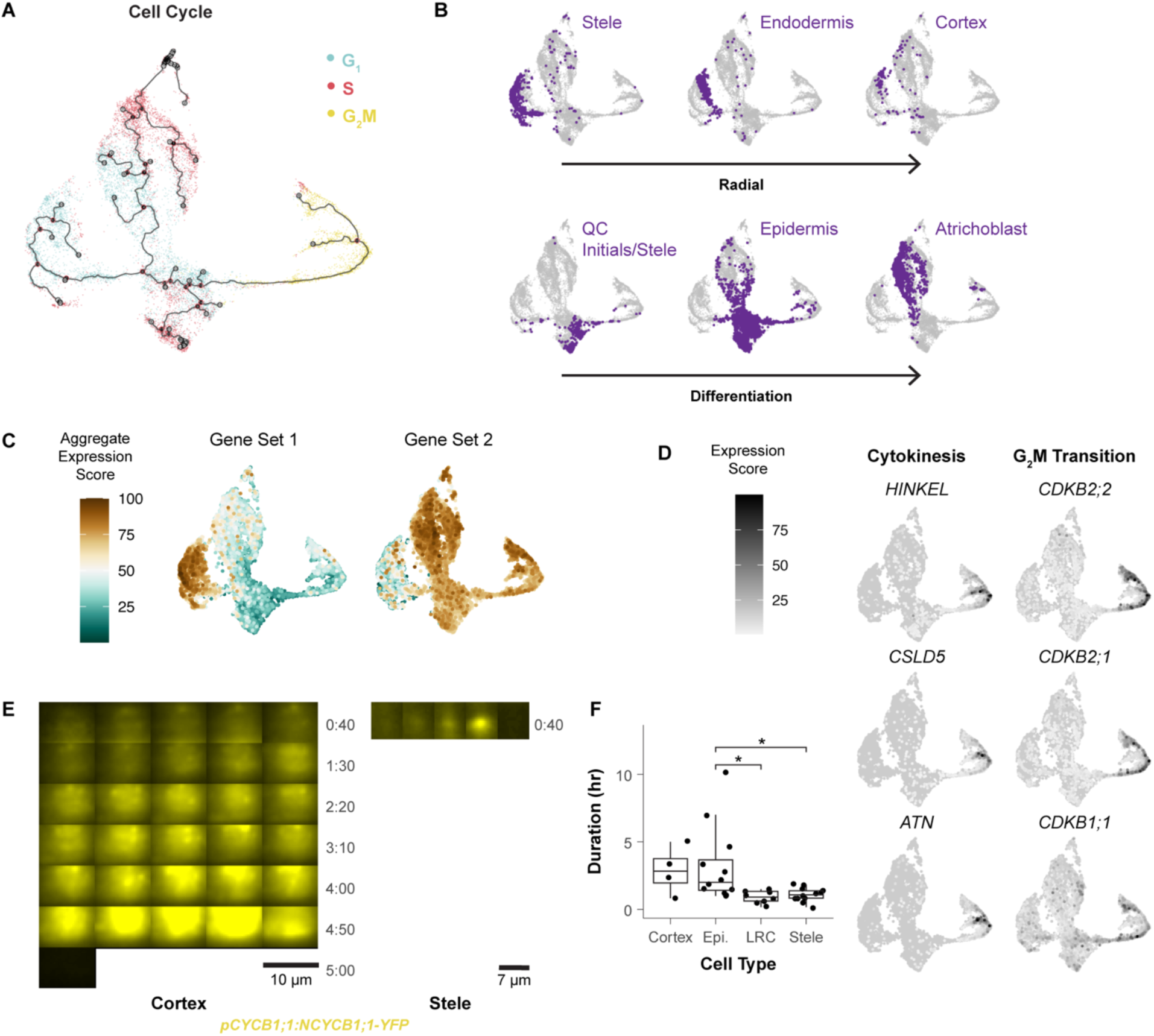
Different cell types follow different trajectories through the cell cycle. (A) Pseudotime map of cells clustered using only cell cycle markers and colored by phase assigned in Seurat. (B) In the same UMAP clustering in A, cells were labeled by their independently determined cell identity, showing groupings by both developmental stage (e.g., QC region) and radial cell identity. Arrows indicate inner to outer cell files (top) and differentiation stage from young to older (bottom). (C) Aggregate expression of genes enriched in either the epidermal or stele-endodermis-cortex G1 branch, which showed genes with differential functions. (D) UMAPs showing the expression of genes specific to sub-regions of the G2/M branch, with representative genes involved in cytokinesis (lower branch) and the G2/M transition (upper branch). (E) Variable lengths of G2/M phase marker expression shown between one cortex and one stele cell. Each frame of each montage is ten minutes apart from time lapse movies taken with a Tilt light sheet system (Mizar). Each montage shows the frame at which *pCYCB1;1:NCYCB1;1-YFP* first becomes visible in the given cell to the frame where the reporter disappears, indicating the end of G2M. (F) Quantification of G2/M duration based on the amount of time *pCYCB1;1:NCYCB1;1-YFP* was visible for in many cells from two time lapses (n=36 cells). Asterisks represent significant differences in G2/M duration (p < 0.05, pairwise t-test). Each dot represents a cell. See also Figure S6, Table S3, and movie S1.

The most apparent trend was a difference in G1 to S branches between inner and outer cell types. Xylem and phloem occupied successive layers of a left branch, with endodermis and cortex on an outermost layer of the branch (Figure 2B). Epidermal cell types occupied a distinct second branch, and a third branch contained the slow cycling cells around the quiescent center (QC), the core of the stem cell niche (Figure 2B). This distinct stem cell behavior is in accordance with the well-documented slower rate of division of these cells compared to the more proximally located (shootward) transit amplifying cells^6,7,45^. The epidermal G1 branch was enriched for genes related to translation, while the stele-endodermis-cortex G1 branch was characterized by gene expression related to cell wall synthesis (Figure 2C, Table S4). This suggests that the specialized functions of specific cell types are at least partially regulated within the cell cycle as they mature in the meristem. The overall patterns were consistent with the hypothesis that plant cells have multiple G1 modes^46^.

In addition, we observed two subpopulations of cells in G2/M, separated into an upper and lower branch (Figure 2A). While many genes were commonly expressed across G2/M cells, we found unique functional enrichments between these branches. The upper branch expressed genes that regulate the G2/M transition, while the lower branch expressed cytokinesis regulators (Figure 2D). These two branches revealed a set of *in vivo* G2/M processes shared by all cell types. To further address cell type specificity of cell cycle markers, we performed an additional differential expression analysis by isolating individual cell types and testing for expression differences between phases. Using these analyses, we generated a matrix of cell type specific phase markers (Table S6).

Nonetheless, we observed cell type-specific biases within the different regions of the G2/M branch, potentially indicating differences in the amount of time cells spent in a given sub-phase of G2/M (phase dwelling). To test this hypothesis *in vivo*, we generated long-term time-lapse light sheet microscopy movies of roots expressing the three-color cell cycle translational reporter, PlaCCI, which marks G1 (CDT1a, cyan), S (HTR13, red), and Late G2 through M phase (CYCB1;1, yellow^18^). We measured G2/M duration in epidermal, cortical, stele, and lateral root cap cells (Figure 2E, 2F, movies S1). From one such time lapse, epidermal and cortical cells remained in G2/M twice as long as stele and lateral root cap cells. But there was also significant variation in G2/M duration within cell types. For example, the length G2/M ranged from 1:00 to 10:10 (hr:min) in the epidermis and 0:10 to 01:50 in the stele (Figure 2F). Thus, live imaging corroborated the cell type-specific phase-dwelling variations detected by the cell cycle mapping of scRNA-seq profiles. Overall, these observations reveal the extent to which the cell cycle is tailored to cell identity and developmental stage.

### Tissue-wide coordinated G1 exit and rapid G1 is linked to regeneration efficiency

Many questions in plant and animal regeneration concern how cell cycle regulation mediates cellular reprogramming. For example, we have observed that cell cycle speed increases during RAM regeneration^47^, but it remains unclear whether this is due to a uniform increase in cell cycle speed across phases or whether certain phases are truncated to achieve fast divisions. Thus, we applied the cell cycle marker analysis to single-cell analysis of regenerating cells in the excised root tip over a time course of 4 to 36 hr post-cut^42^ (hpc) to analyze their fine-grained cell cycle dynamics. We focused on the relatively small set of cells actively contributing to regeneration, as previously annotated^42^, aligning regenerating single-cell profiles in cell cycle pseudotime, similar to above. The analysis showed that regenerating cells disproportionately accumulate at the G1 to S transition and are largely absent from G1 phase (Figure 3A).

**Figure 3:**
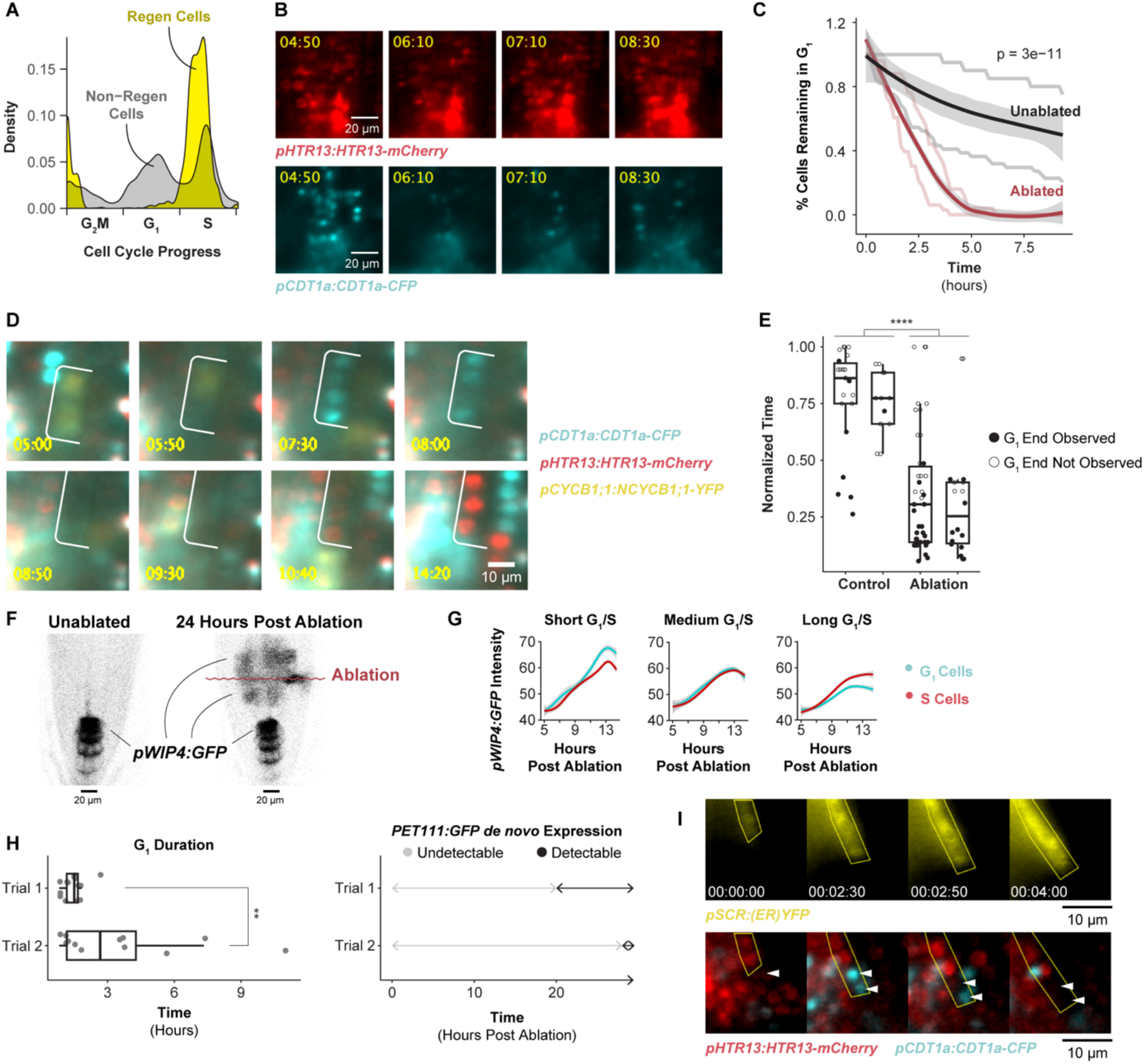
The G1 phase of the cell cycle is dramatically truncated in regenerating cells. (A) Summary of the frequency of a given cell cycle phase in regenerating (yellow) and non-regenerating (grey) cells. Cells are aligned along a cell cycle pseudotime on the x axis, with their density shown on the y axis with G1 predominant in non-regenerating cells and almost absent in regenerating cells. (B) Representative images of cells coordinately exiting G1 following tissue damage (bottom). S-phase cells (top) serve as a control showing a continuous strong signal (no depletion) in the same roots. Time is shown in hours:minutes and 00:00 is the time of the ablation. (C) Quantification of the coordinated G1 exit shown as a survival analysis of the population of G1 cells identified at the beginning of the time lapse. The y axis represents the fraction of G1 cells from time zero (the first frame of the time lapse) still in G1 at time n. The x axis shows time since the beginning of the time lapse. Time zero for each ablated root is 3 hpa. Two unablated and two ablated roots were analyzed. (n = 165 cells, p-value = 3e-11, log rank test). Lightly colored lines are individual replicates, and the bold line is a LOESS regression of the two trials. P-value of survival rates is significantly different between ablated and unablated roots using the log rank test. (D) Representative time-lapse series of a short G1 in an ablated root in which cells pass through the phase in as little as 2 hr. Time is shown in hours:minutes and 00:00 is the time of the ablation. (E) Quantification of G1 duration in control and ablated roots for two trials. Time is normalized within each root to the duration of the time lapse in which it was measured, where the y-axis represents the time in which a cell was in G1 divided by the duration of the time lapse. Filled dots represent cells in which the end of G1 was observed. Each dot represents an individual cell. (n=94 cells, p-value = 3.221e-09, Mann-Whitney U Test). (F) Representative image of the *pWIP4:GFP* expression domain before ablation and 24 hpa. The red wavy line marks the location of the ablation. (G) Quantification of *pWIP4:GFP* signal over time in G1 and S phase cells, with different plots showing analysis of cells grouped by the length of G1 or S. Short = 3 hr, medium = 6 hr, long = 9 hr. (n = 650 cells in 1 root). (H) Quantification of G1 duration in two roots (left) and the timing of *PET111* expression establishment in the regeneration zone in the same two roots (right) showing the association between G1 duration and PET11 appearance. Each dot represents an individual cell. Trials refer to individual root time lapses (p-value = 0.00413 Mann-Whitney test). (I) Representative image of the expansion of the expression domain of the *pSCR:erYFP* reporter during regeneration. The SCR expression domain is outlined with a yellow ROI on both the upper and lower panels. Panels were chosen to show the cell marked with an arrow just before division, while in G1, and after each daughter exits G1. The first frame of this montage is 26 hpa and the time stamp of each frame are as follows (day:hr:min): 00:00:00, 00:02:30, 00:02:50, 00:04:00. See also Figure S7, Table S4-S5s, and Movies S2-S3.

This result suggests that G1 is dramatically shortened relative to the other phases of the cell cycle during early regeneration, raising the possibility G1 is the phase that is dramatically altered during regeneration. To measure G1 duration together with fate re-specification, we used time-lapse light sheet imaging on live regenerating roots, quantifying G1 phase duration concurrently with cell fate changes. In the root tip regeneration system, the meristem is excised, completely removing the QC and columella cells, which are then respecified within a day from vascular and other cells left in the cut stump^2^. To enable rapid imaging after regeneration, we generated a similar root-tip excision using a two-photon ablation system in which the root meristem is essentially isolated by a plane of dead cells causing regeneration of QC and columella shootward, as in root tip excision (see below). We used the PlaCCI^18^ marker to track cell cycle phase and the QC-columella marker *pWIP4:GFP* to track cellular reprogramming^48^. Return of WIP4 expression shootward of the ablation site marks cells that are in the process of adopting QC and columella fates in the newly formed meristem – a key step of RAM regeneration. By monitoring this region, we could track the full history of cell cycle phases, their duration, and reprogramming state.

We observed that cells in the regeneration zone coordinately exited G1 approximately 6 hours post ablation (hpa), within 1 to 2 hr of one another, depending on biological replicate, and prior to new *pWIP4:GFP* expression (Figures 3B and 3C, Movies S2, S3). We formally tested whether this behavior differed from G1 cells in unablated roots with a survival analysis by identifying G1 cells in the first frame of a given time lapse and tracking how long it takes that population of cells to transition to S phase. We observed a gradual population decay in unablated roots, and a much more rapid decay in ablated roots (log rank test p-value = 3e-11). This is consistent with the dramatic depletion of G1 cells detected in the scRNA-seq analysis of regenerating cells.

Following this coordinated G1 exit and between 8 to 12 hpa, these cells then proceeded through G1 at an accelerated rate (Figure 3D, 3E). To quantify G1 length, we measured the elapsed time between when CDT1a became visible after mitosis (early G1) to when CDT1a was degraded, indicating S-phase entry. Some of the observed G1 events did not end during the time lapse in both the control (76 percent) and the ablation (38 percent) movies. In these cases, we measured G1 duration in three ways: 1) as the time between when CDT1a became visible and the final frame of the time lapse, 2) as equal to the observed G1 duration time for this region of the root, which is estimated to be longer than 20 hr^7^, and 3) as the fraction of total movie duration (Table S5). By all these metrics, the difference in G1 duration is statistically significant (Mann-Whitney test, p-value = 1.614e-08, p-value = 2.04e-05, or p-value = 3.221e-09). Thus, the specific, highly localized set of cells that will reprogram to generate the new root tip undergo much more rapid G1 than their neighbors. Overall, the data shows there are two G1 regulation phenomena apparent in these time lapses: coordinated exit (Figure 3B, 3C) and subsequent rapid progression (Figure 3D, 3E).

To test the association between rapid G1 and reprogramming, we identified cells that eventually expressed the *pWIP4:GFP* marker (indicating cellular reprogramming, Figure 3F) and analyzed their cell cycle dynamics retrospectively in the time-lapse movies. We compared the timing of re-specification in cells with short G1 vs. neighboring cells that displayed longer G1s (Figure 3G). We categorized cells based on short, medium, and long G1 and S duration. While cells gained *WIP4* marker expression at a similar rate regardless of S phase duration, cells with short G1 gained higher *WIP4* expression levels than nearby cells with long G1 (Figure 3G). There was no relationship between *WIP4* expression and G1 duration in unablated roots (Figure S7). Thus, a specific group of cells in the regenerating stump that undergo fast G1 reprogram more rapidly than slower G1 neighbors.

To determine whether the relationship between G1 length and re-specification holds for other markers that are expressed later during regeneration, we looked at a late-stage marker for columella, *PET111:YFP*. In this case, we exploited variability in *PET111:YFP* return time and G1 duration between roots to explore whether these two variables were correlated. In this analysis, G1 duration was broadly predictive of PET111 re-appearance (Figure 3H). For example, a root in which the median G1 duration was 1.5 hr began to express *PET111:YFP* in the regeneration domain at 20 hpa. A second root in which median G1 duration was 2.7 hr began to express *PET111:YFP* at 28 hpa.

Having associated the return of two distal cell fate markers, *WIP4* and *PET111*, with G1 duration, we sought to test a third marker of another cell type. We chose to observe endodermal fate establishment late during regeneration with the *pSCR:erYFP* reporter in the PlaCCI background. New endodermal fate establishment is a rare event, and across two time lapses we observed five cases of cells establishing *de novo* SCR expression. In each of the five cases, *de novo* expression was established while cells were in G1 phase of the cells cycle (Figure 3I). Thus, rapid G1 phases in plant root regeneration are tightly associated with reprogramming of excised cell fates. This opens the possibility that rapid G1 could play a functional role in promoting cellular reprogramming in plants.

### GSH is enriched in G1 nuclei at steady state and immediately following tissue damage

Having implicated G1 duration in control of regeneration efficiency, we next sought to establish a mechanistic link between injury and cell cycle regulation. The finding above showing “response to wounding” and “response to oxygen-containing compound” terms enriched in G1 was intriguing because ROS has potential links to both the cell cycle and wounding (Figure 1F). In particular, GSH is the primary antioxidant in the cell, and GSH levels in the nucleus have been found to vary over the course of the cell cycle in both plants and animals^32^. In Arabidopsis, GSH has been demonstrated to be necessary for the G1 to S transition in root formation^30^. In addition, ROS generation is a hallmark of tissue damage^35^, with variants in genes controlling thioredoxin-mediated ROS associated with natural variation in regeneration capacity in Arabidopsis^49^. Thus, we reasoned that G1 cells could be primed to respond to ROS signals generated by tissue damage.

To explore this connection, we performed live imaging with the ROS indicator H2DCFDA and the GSH dyes Blue CMAC and CMFDA during stereotypical root growth and regeneration (Figures S8 and S9). We first confirmed that these dyes had no effect on meristem size and regeneration efficiency (Figure 4A). We used time-lapse confocal imaging and the ablation described above to observe GSH localization within the first 30 minutes of tissue damage. We distinguished cells in G1 vs S phase using the PlaCCI marker (noting that the S-phase mCherry marker is also expressed in early G2 phase^18^). We segmented nuclei based on the S-phase mCherry marker of PlaCCI and found that, in control roots, Blue CMAC signal was higher in G1-phase nuclei than in S phase nuclei (Figure 4B), building on prior evidence that suggested nuclear GSH controls the G1 to S transition^30,33^. In regeneration, we observed a pulse of nuclear GSH immediately after ablation just above the injury site in segmented nuclei across all cell types (Figures 4C-4E, Movie S4). In addition, in the root cutting injury, at the 2- and 4-hours post cut (hpc) time points, nuclei that showed the highest CMAC signal shootward of the cut site were in the same region in which cells undergo rapid G1 phases (Figure S10), and GSH remains high in G1 cells through 24 hpc (Figure 4F). Overall, the results suggested that the earliest cells to reprogram first undergo a local burst of GSH import into the nucleus then exhibit a coordinated G1 exit followed by a rapid G1 phase.

**Figure 4:**
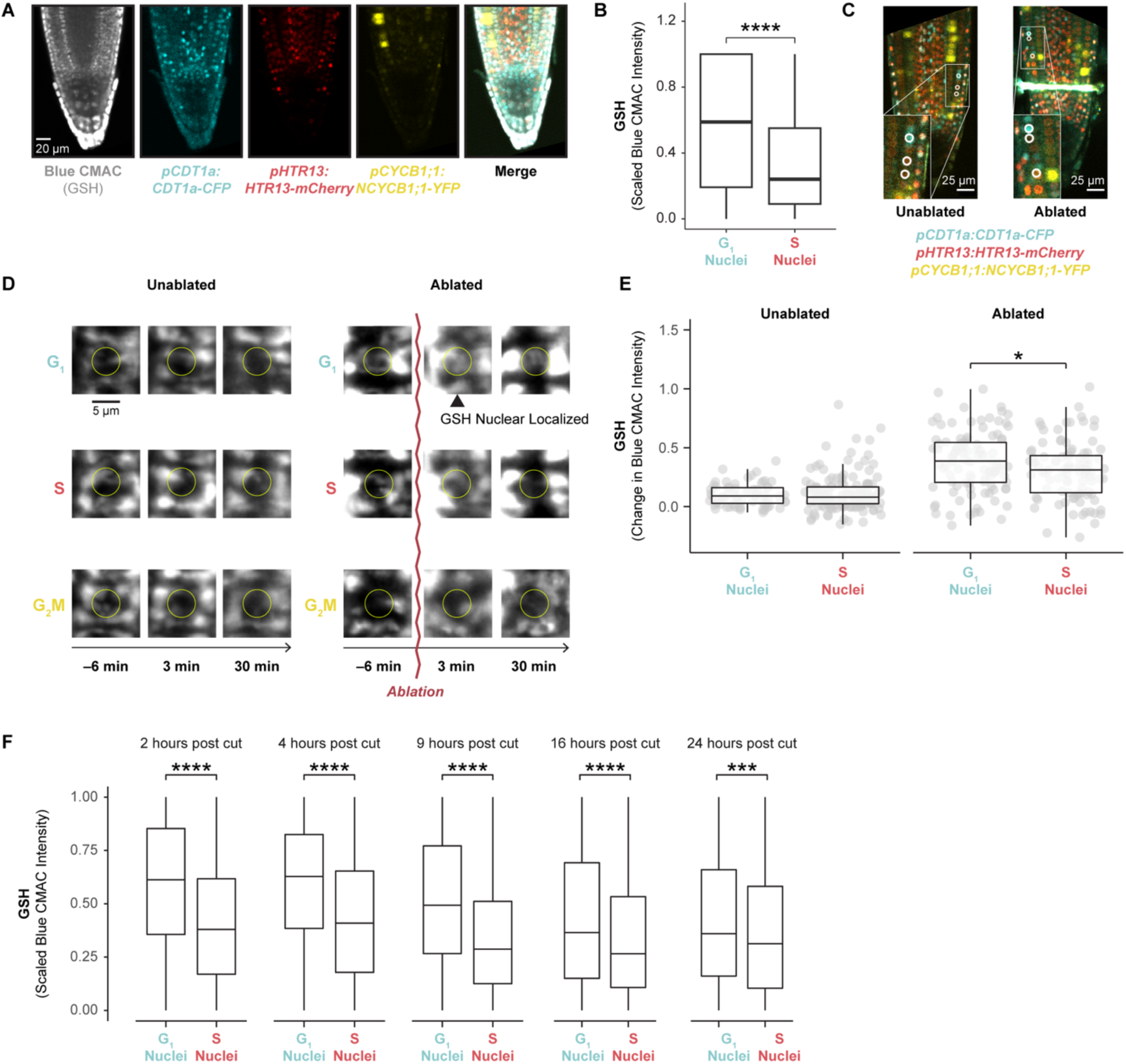
Regenerating cells import glutathione to the nucleus in G1. (A) Representative confocal microscopy image of a PlaCCI seedling stained with Blue CMAC overnight, showing phase markers and Blue CMAC staining. (B) Quantification of Blue CMAC in G1 and S phase nuclei. (n= 416 nuclei, n=2 roots, p-value = 2.23e-10, student’s t test). (C) Images showing the location of cells analyzed in 4D annotated (circles) and shown in insets. All cells in these images were analyzed in 4E. (D) Representative images of cells in each phase of the cell cycle in control and ablated roots shown in a time-series montage. Time relative to ablation is shown. The yellow circle shows the position of the nucleus. (E) Quantification of the change in Blue CMAC levels in nuclei of cells in G1 and S phase in unablated and ablated roots. Each boxplot shows GSH signal measured in segmented nuclei from frame 3 of the relevant time lapse. In the case of the ablated root, this frame was taken 3 minutes post ablation. The y-axis represents the amount of nuclear GSH signal at frame three relative to the amount of GSH signal in those same nuclei at frame 1. Each dot represents a nucleus. (n=420 nuclei and 2 roots, p-value = 0.0296, student’s t test). (F) GSH levels are shown in segmented nuclei at various time points post cut in G1 and S phase cells (n=8632 nuclei, n=35 roots, *** <5e-4, ****<7e-9,student’s t test). See also Figure S8-S10 and Movie S4.

### GSH depletion inhibits regeneration efficiency

To explore the functional role of GSH in regeneration, we depleted GSH during regeneration using the GSH synthesis inhibitor, L-Buthionine-sulfoximine (BSO), following established protocols^31^. We first depleted GSH in roots by germinating seedlings on plates supplemented with 1 mM mM BSO (Figures 5A and 5B). We performed this experiment on seedlings expressing PlaCCI and the *pWIP4:GFP* transcriptional reporter to simultaneously track cell division and QC reestablishment, while also using Blue CMAC staining to visualize GSH.

**Figure 5:**
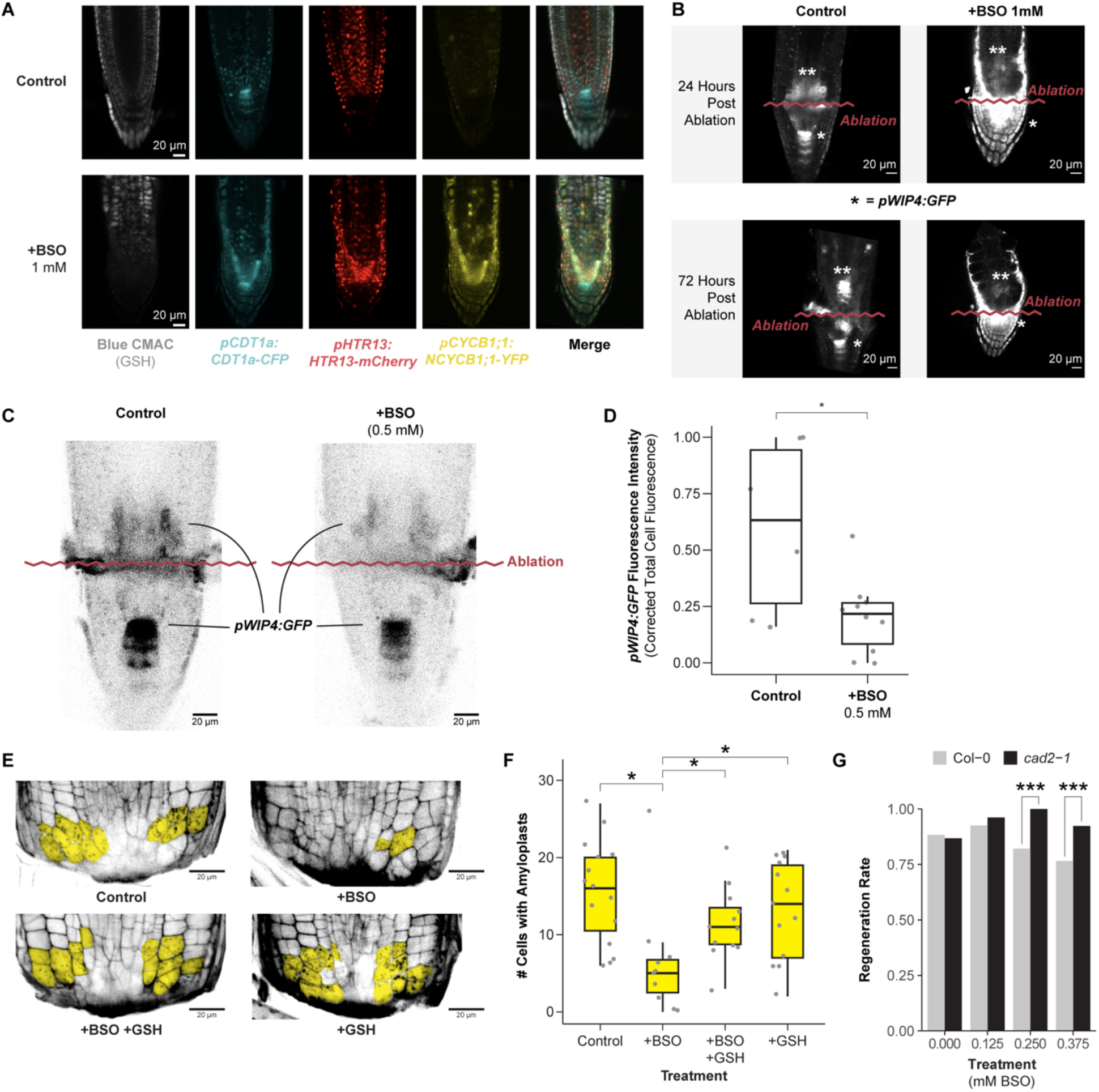
Depletion of GSH biosynthesis with BSO impairs regeneration and is rescued by exogenous GSH. (A) 7 days post germination (dpg) root meristems (PlaCCI x *pWIP4:GFP*) grown on MS (control) or on MS+1mM BSO then stained overnight with Blue CMAC. (B) Representative images of the *pWIP4:GFP* signal in a median section of a control and BSO-treated root at 24 and 72 HPA. The ablation site is indicated with a wavy red line. The original QC within the stem cell niche is indicated with an asterisk (*), and the newly forming stem cell niche is marked with two asterisks (**). (C) Representative images of *pWIP4:GFP* signal 24 hpa in control and 0.5 mM BSO treatment. (D) Quantification of *pWIP4:GFP* signal in the regeneration zone of roots 24 HPA in control and 0.5 mM BSO treatment. The y-axis is the corrected total cell fluorescence of *pWIP4:GFP* in the new QC domain scaled to render experiments comparable between technical replicates (n=16 roots, p = 0.05 Wilcoxon test). (E) Representative images of regenerating root tips stained with mPS-PI to visualize cell walls and amyloplasts 18 hpc. Cells with amyloplasts are pseudo-colored in yellow. The treatments are control, 0.5 mM BSO, 0.5 mM GSH, or combined 0.5 mM BSO + 0.5 mM GSH. (F) Quantification of the number of cells with amyloplasts in a population of roots from each treatment group shown in E. (n=48 roots, * < 0.03, Wilcoxon test). Each dot represents a root. (G) Root tip regeneration rates (y-axis) for col-0 (grey) and *cad2-1* (black) seedlings grown on increasing concentrations of BSO (x-axis). At higher concentrations, col-0 regenerates significantly less efficiently than *cad2-1* (n>65 for each treatment group, p-value < 0.003,chi square test).

The control seedlings regenerated a new QC shootward of the ablation over the course of 72 hr (Figure 5B). Seedlings germinated on 1mM BSO were depleted for Blue CMAC signal (Figure 5A), and they showed weak *pWIP4:GFP* expression shootward of the ablation through 72 post injury. In addition, these roots failed to form an expression pattern indicative of new QC establishment (Figure 5B). However, as previously shown^31^, we observed that most seedlings treated in this manner had short roots before the ablation, raising the possibility that meristem defects before ablation impaired regeneration. To address this issue, roots were germinated on a lower concentration of BSO (0.5mM) on which they displayed normal morphology^31^. Although ablated roots grown on this lower BSO concentration eventually regenerated, they showed a lower amount of *pWIP4:GFP* expression in the regeneration zone at 24 hpa (Figures 5C and 5D). Thus, depletion of GSH to a level that does not affect stereotypical root growth still impairs the re-specification of the columella and QC marker.

Columella cells are necessary for the root to sense gravity, which requires ballast-like organelles called amyloplasts that settle along the gravity vector. Thus, amyloplasts are a functional marker for columella re-specification. To quantitatively assess the effect of GSH depletion on regeneration efficiency, we monitored the number of cells containing amyloplasts in excised root tips with modified Pseudo Schiff-Propidium Iodide (mPS-PI) staining^50^ at 18 hpa in four conditions: mock, 0.5 mM BSO, 0.5 mM GSH, and 0.5 mM BSO + 0.5 mM GSH combined. We found that treatment with BSO significantly decreased the number of cells with *de novo* amyloplasts at 18 hr and that co-treatment with GSH rescued regeneration to the level of untreated roots (Figures 5E and 5F), consistent with regeneration defects caused by diminished levels of GSH post-injury. Finally, in order to confirm that BSO inhibits regeneration specifically by depleting GSH, we performed gravitropism experiments with *cad2-1* roots on increasing doses of BSO (Figure 5G). This mutant line has a point mutation in the domain where BSO physically interacts with GSH1^51^, rendering these seedlings insensitive to BSO treatment. At higher concentrations, 0.25 mM and 0.375 mM, *cad2-1* roots regenerated more efficiently than wild type, indicating that BSO specifically inhibits regeneration via its effect on GSH biosynthesis.

We next directly tested whether BSO treatment perturbs G1 dynamics during regeneration by performing long-term time-lapse imaging in the PlaCCI line with roots germinated on 0.5 mM BSO. We found that following injury, G1 cells in BSO-treated roots failed to undergo the coordinated exit that we observed in untreated roots (Figure 6A). We formalized this observation by performing a survival analysis, recording how long it took cells in G1 at the beginning of the time lapse to enter S phase (Figure 6B). We found that the time cells remained in G1 was significantly prolonged in BSO-treated roots (log rank test; p-value < 2e-16). Interestingly, BSO appeared to have a greater effect on cells away from the immediate injury site. For example, in BSO-treated roots, most cells above the injury failed to exit G1 in a coordinated manner, while the first two or so layers of cells near the cut site still showed the coordinated exit despite BSO treatment (Figure 6D). This is consistent with a gradient of GSH that is highest in cells immediately adjacent to the wound site dissipating in more proximal cells, where BSO was presumably more competent to disrupt GSH signaling.

**Figure 6:**
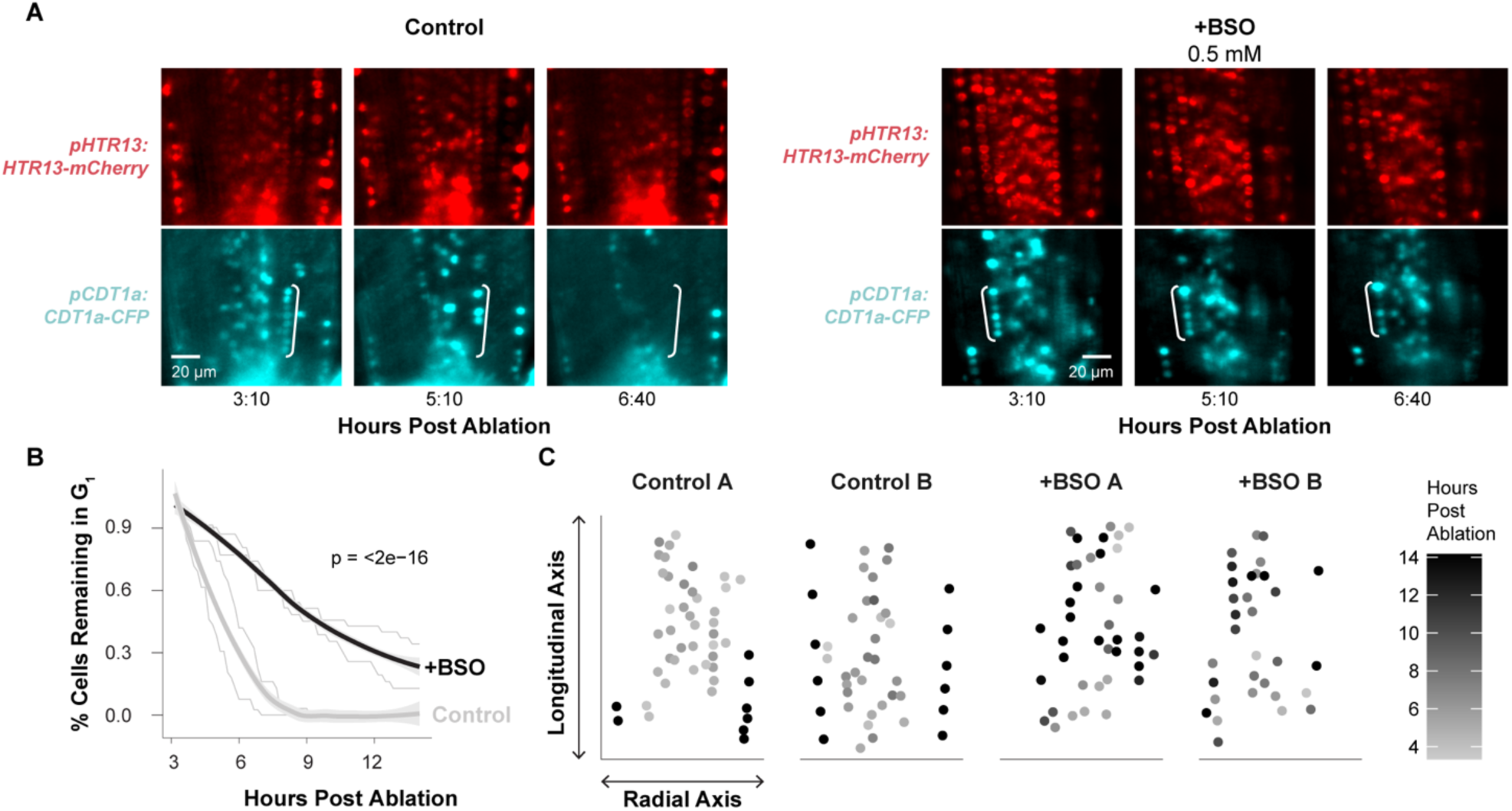
Depletion of GSH with BSO eliminates the coordinated exit from G1 and increases G1 duration in regeneration. (A) Representative images from a control (left) and BSO treated time lapse (right) immediately shootward of the ablation site, showing the S phase (red) and G1 (cyan) markers. Cells from the cortex in G1 are bracketed to highlight the differential disappearance of G1 phase cells in control vs. post ablation. S phase cells are shown to confirm no change in their fluorescent signal. (B) G1 duration quantified in survivor curves, where cells in G1 were identified in the first frame of the time lapse and tracked until their transition to S phase for control (gray line) and BSO-treated (black line) time lapse experiments. Lightly colored lines are individual replicates, and the bold line is a LOESS regression of the two trials (p=<2e-16, n = 126 cells, controls are the same cells shown as “Ablated” in Fig. 3C, log rank test). (C) Grayscale representation of the time in hpa that cells exit G1 mapped onto the given cell’s coordinates within the roots, where the Y-intercept represents the ablation site and each dot represents a cell, with two example roots per condition (A,B). Shading scale represents time post ablation when a cell exited G1 phase.

Overall, the effects of the BSO treatment on *pWIP4:GFP* expression levels (Figure 5C, 5D), amyloplast formation (Figures 5E and 5F), and G1 dynamics lead to the conclusion that GSH depletion slows regeneration at least in part through modifying G1 exit and duration.

### Ground tissue is an apparent source of glutathione in growth and regeneration

In our staining for GSH in unablated roots, we observed a striking pattern in which Blue CMAC was highly localized to the cap, epidermis, and ground tissue (cortex and endodermis), while the stele stained much more weakly (Figure 4A, leftmost panel). The pattern did not appear to be an artifact of limited cell penetration, as the two GSH dyes Blue CMAC and CMFDA have similar staining patterns, while the ROS indicator H2DCFDA, which has a similar chemical structure to CMFDA^52^, stains all files relatively evenly (Figure S8A). In particular, with both Blue CMAC and CMFDA, we observed highly concentrated staining in the endodermis and cortex (Figure S8A). The localization pattern was consistent with independent data we gathered from scRNA-seq studies that showed GSH biosynthesis genes are also highly expressed in the ground tissue (Figure S8B). This led us to hypothesize that ground tissue could be a source of GSH for root growth and rapid dissemination upon injury.

Metabolites and other small molecules can travel rapidly between plant cells through symplastic connections that form tunnels between adjacent cell walls called plasmodesmata^53^. To ask whether ground tissue could serve as a source of GSH for other files to enable homeostatic growth and regeneration, we employed a callose synthase induction system that blocks symplastic transport out of endodermis and the cortex with an estradiol-inducible callose synthase driven using the *J0571* enhancer trap^54^ and then assayed for growth (Figure 7A) and regeneration efficiency (Figure 7B). Exogenous GSH is known to enhance growth rates in Arabidopsis roots, so we controlled for the nonspecific effects on growth by comparison to high sucrose (1% versus the standard 0.5%), which also enhances root growth. Accordingly, both sucrose and GSH both increased growth rates in control roots. However, only GSH-treated roots partially rescued the growth of the ground-tissue blocked roots (Figure 7A). Furthermore, after injury and symplastic block of the ground tissue, GSH, but not sucrose, rescued regeneration efficiency (Figure 7B). Finally, we confirmed induction of callose synthase in the ground tissue resulted in depleted GSH in the stele by staining induced roots with Blue CMAC (Figure 7C). We quantified Blue CMAC signal in the stele in induced and uninduced roots and found signal was statistically significantly decreased in the stele but not in other cell files (stats n=21 roots, p-value = 0.0041). The results suggest that ground tissue is a source of GSH for normal growth and tissue regeneration, licensing rapid exit from G1, an abbreviated cell cycle, and rapid cellular reprogramming.

**Figure 7:**
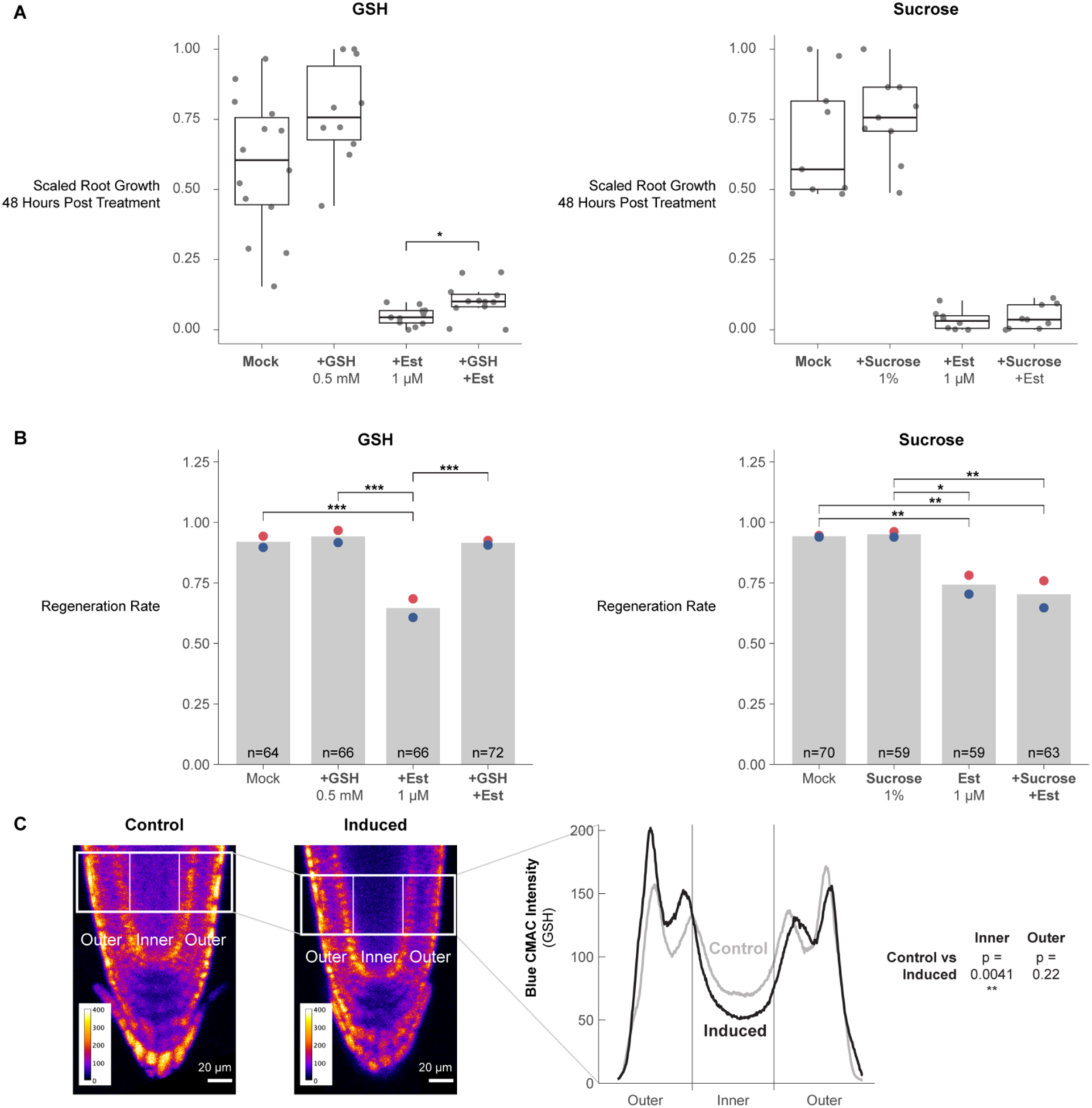
The ground tissue is an apparent source of GSH in homeostatic growth and regeneration. (A) Root growth (y-axis) post callose-synthase induction for each treatment condition (x-axis). Root lengths are scaled to their own controls within technical replicates from 0 to 1 to render them comparable across batches. Statistical significance was determined by the pairwise Wilcoxon test comparing estradiol or non-estradiol categories (i.e. mock was tested versus GSH and estradiol was tested versus estradiol + GSH) (n > 10 roots per condition, p-value = 0.02 by the Wilcoxon test). Each dot represents an individual root. (B) At left, regeneration rates (y-axis) based on the gravitropism test at 48 hpc. The conditions (x-axis) are control (mock), GSH treated roots (+GSH), estradiol-treated roots (+Est, induction of callose synthase expression to block transport out of the cortex and endodermis), estradiol + GSH treated roots (+GSH,+Est). At right, the same treatments substituting 1 µM sucrose for GSH. Red and blue dots represent the regeneration rates of technical replicates (***<0.00071, **<0.0003, *<0.004,Fisher’s exact test). (C) In the left panel, representative confocal microscopy images of GSH staining using Blue CMAC for uninduced control (left) and ground tissue callose-synthase induced (right) roots are shown. ROIs on the images show representative examples of ROIs used to calculate Blue CMAC intensity on the right panel across inner and outer files. The y-axis on the right panel represents the average intensity for each column of pixels of several comparable ROIs (n=21) across the x dimension of the ROI. Average intensities for inner versus outer ROIs were tested for significant difference with, with a significant difference only in the inner cell files (p=0.0041, pairwise t test).

## Discussion

There are clear connections between cell division and cell fate decisions across the kingdoms of life. In the context of root tip regeneration, cell division is known to be necessary to enable complete root tip repair after excision^2^. Here, we leveraged the ability to induce cellular reprogramming and closely monitor cells with both time-lapse microscopy and transcriptomics in root tip regeneration to demonstrate that the rate of cell division, mediated by dramatic alteration of G1 phase, has a direct influence on cellular reprogramming. In our findings, glutathione (GSH) mediates fast divisions via truncated G1 phase in a small number of cells that will go on to reprogram their fate first. Furthermore, we found that the ground tissue is an important source of glutathione for stereotypical growth and regeneration. Given the findings, we posit a model in which GSH produced in the ground tissue rapidly disseminates to nearby tissues after injury. The GSH stores are preferentially imported into the nucleus of a subset of cells near the injury site where they instigate coordinated exit from G1 and accelerated cell divisions that permit rapid cellular reprogramming.

### The root has distinct inner, outer and promeristem cell cycles

Using bulk and scRNA-seq, we defined a novel list of cell cycle phase markers, including a large set of G1 markers, which now provide a resource for the plant community, particularly for analysis of the cell cycle in scRNA-seq studies. By clustering cells from an unsynchronized dataset^42^ based on cell cycle-regulated genes, we detected multiple paths through G1 indicating that distinct G1 states exist in Arabidopsis roots (Figure 2A). We took several steps to ensure that cell type-specific markers were filtered out of the cell cycle phase markers, even if they appeared to be phase-specific. Nonetheless, mapping cell identities onto the cell cycle trajectory revealed that different cell types appear to prefer unique paths through the cell cycle (Figure 2B, Figure S6). For example, in one G1 phase pseudotime path, xylem and phloem cells appeared to occupy distinctly different layers of a left branch, while trichoblasts largely occupied a central branch. Genes expressed in the xylem-phloem branch are enriched for GO terms relating to auxin response and developmental processes (Figure S6). The genes that were selected as cell cycle phase markers are widely expressed across cell types, so the groupings by cell identity must reflect how these commonly expressed genes are specifically regulated in a given cell type. It is feasible that common facets of plant cell biology--such as construction of a cell wall, which also varies among cell types^55^—are linked to changes in the cell cycle^56^ to accommodate differences among cell types.

The most robust cell-cycle markers we identified represent non-canonical cell cycle genes (Figure 2E). This is evidenced by the fact that, while core cell cycle regulators behave well in our datasets (Figure 1C, 1D, Figure S5), few genes we identified as being most highly cell cycle-regulated are among the genes considered to be core cell-cycle regulators (Table S1). It has been argued previously that different occurrences of cellular quiescence in plants – meristematic quiescence, dormancy, and terminal differentiation – are regulated distinctly and by non-canonical cell cycle genes^46^. Our results show that multiple subpopulations of G1 cells exist and that they are characterized by the expression of distinct transcriptional modules. One subpopulation is characterized by the expression of genes relating to cell wall synthesis, while the other is characterized by genes regulating translation, both of which are functions that have been tied to the G1 phase in plants^56,57^. Another recent report has shown that the longitudinal axis of the root is largely due to variation in G1 length^7^. Our results support a general model in which the cell cycle is finely tuned to both the maturation stage, as is well documented, but also to cell identity.

In addition to the ability to detect multiple G1 phase cell populations, we also find evidence for two G2/M populations, which express genes related to checkpoint regulation or cytokinesis respectively. In parallel, our *in vivo* data shows that distinct cell types spend different amounts of time in G2/M in the RAM. This indicates our cell cycle marker set can be used to detect cell cycle sub-phases in Arabidopsis scRNA-seq data and enable further dissection on cell cycle regulation in existing and future plant scRNA-seq datasets.

### Reprogramming plant cells divide rapidly by shortening G1

In metazoans, evidence links rapid G1 phases with competence to reprogram^58,59^. For example, embryonic stem- and induced-pluripotent cells are characterized by rapid cell cycles with short G1 phases^58^. In plants, while division times in the indeterminately growing meristems are about 20 hr^6^, cell division rates during embryogenesis, lateral root formation, and root regeneration – all instances of novel root formation rather than homeostatic growth – show a dramatic acceleration to 3 to 7 hr^23,24,47^. Here, we show that the fast divisions in regeneration are largely due to a highly truncated G1, consistent with data from efficiently reprogramming murine hematopoietic progenitor cells^59^--another context in which dramatic changes in the cell cycle are mediated by alterations in G1.

G1 has been shown in metazoans to be a key point in which cells are receptive to signals that promote specialized cell fate and differentiation^60–62^. Thus, it has been posited that rapid G1s allow cells to remain pluripotent by avoiding differentiation signals^60–62^. In our scRNA-seq profiles, we did not detect any enrichment of known cell identity markers in a given phase of the cell cycle. Thus, we have no evidence that short G1s could bypass differentiation signals, although we cannot rule out that cell fate markers are induced synchronously but transcribed at different rates or regulated at another level as has been shown for some specific contexts in plants^3,4^. Nonetheless, our experiments clearly associate rapid G1 phases and coordinated G1 exit with the competence to reprogram cell fate across cell types. Importantly, neighboring cells that did not undergo rapid G1 phases could still reprogram, and, while treatments that perturbed G1 coordination showed slower regeneration dynamics, even injured roots exposed to such treatment eventually regenerated. Thus, rapid and coordinated G1s are not absolutely necessary for cellular reprogramming. It is not clear if G1 dynamics during regeneration have a direct role in avoidance of differentiation signals, or, if rapid G1s might simply allow a faster entry into S phase. While mechanisms have been identified to link maintenance of histone modifications to DNA replication in plants^63,64^, there is inherent potential for remodeling the chromatin landscape during DNA synthesis through new histone deposition^65^.

Another possibility is that regulation of G1 may simply be the best option to control overall speed of the cell cycle. Fast G1 phases might not promote reprogramming per se, but could be correlated with other processes that do facilitate reprograming. Several studies have shown that wound responses in plants reflect a bet-hedging strategy that balances defense responses with regenerative growth^66–69^. A similar bet-hedging strategy may have evolved to control cell cycle speed. It has been observed that plant stem cells divide infrequently to limit accumulation of replication-induced mutations^70^. However, wounding creates stresses, such as increased susceptibility of plants to pathogens^71^, that require a rapid response. An ability to trigger fast divisions in otherwise slow-dividing cells may have evolved to limit risks of pathogen exposure following wounding. The ability to pass through G1 quickly and enable rapid divisions may simply represent an adaptation that permits more rapid wound healing and leaves the plant less vulnerable to pathogen attack. Of course, rapid G1s could have multiple roles in regeneration due to a combination of factors.

### G1 cells are primed to perceive tissue damage via GSH nuclear influx

Prior studies have shown evidence that GSH is necessary for the G1 to S transition^30^, while *in vitro* experiments showed that GSH is imported into the nucleus during G1^33^. We showed here that GSH is enriched in G1 nuclei during normal development (Figure 4B) and is transiently increased in G1 nuclei following tissue damage (Figure 4D, 4E, Figure S10). We further present evidence that this transient influx regulates G1 exit during regeneration *in vivo* (Figure 6). Our live-imaging experiments showed that GSH is rapidly nuclear localized in G1 cells, some of which will go on to become the new stem cell niche. This implies that G1 nuclei are inherently more able to take up GSH than those of cells in other phases. Interestingly, when BSO treatment is used to deplete GSH, we find that only the cells closest to the wound site maintain coordinated G1 exit (Figure 6D). This appears to reflect higher levels of GSH closer to the wound, which is feasibly a source of GSH following injury.

Together the evidence leads to a model that could potentially link damage-sensing with cell cycle regulation. G1 nuclei are primed to perceive damage to neighboring cells via GSH nuclear permeability. In this model, GSH released from lysed cells, either directly or by modulating overall nuclear ROS, serves as a damage-sensing signal that allows plant cells to respond to injury by increasing cell cycle speed in close proximity to the wound site.

How G1 nuclei maintain higher GSH permeability than nuclei in other phases of the cell cycle remains an open question. While there is good evidence that the OPT family of genes control intercellular GSH transport in plants (reviewed in^72^) and the CLT family of genes control GSH transport between the cytoplasm and plastids^73^, the mechanism through which GSH is preferentially imported into G1 nuclei in plants is not known^74^. In animals, Bcl-2 has been implicated in the GSH nuclear import^75^. However, plants have no apparent orthologs to Bcl-2. Looking forward, identification of the mechanism responsible for mediating transport of GSH into G1 nuclei will represent a key link between wound signaling and cell cycle regulation in plants.

### Regeneration competence is associated with high levels of GSH across kingdoms

Several lines of evidence in our study pointed to a special role for the endodermis and outer tissues in controlling GSH availability. First, regeneration was impaired when we inhibited the movement of GSH out of the endodermis and cortex (ground tissue) by blocking symplastic connections (Figure 7). Even though both GSH and sucrose enhanced plant growth in general, GSH--but not sucrose--could rescue inhibition of regeneration caused by endodermis and cortex symplastic isolation. In addition, our staining experiments showed GSH is enriched in the endodermis and cortex (Figure S8)--the same tissue where our independent scRNA-seq experiments showed the enzyme for the rate limiting step in GSH synthesis highly enriched^42^.

There is ample evidence that the ground tissue has a specific role in controlling root growth. First, mutants that affect ground-tissue identity, such as *scr* and *shr*, lead to severely stunted roots^76,77^. In addition, it was shown that rescuing SCR function only in the endodermal tissue (leaving out its quiescent center domain) partially rescues *scr* mutants’ growth defect^78^. Some of the endodermal control appears to be mediated by hormone signaling, particularly during stress (reviewed in^79^). Our data suggests that another way that the endodermis controls growth is as a source of GSH to promote G1 exit and advance the cell cycle. In addition, we implicate a unique role for the endodermis in regeneration where it appears to provide a rapid flux of GSH through plasmodesmatal connections (Figure 7).

The association between GSH levels and the competence to regenerate is another trait shared across kingdoms. In animals, the liver also has the highest capacity to regenerate among solid organs^80^. The liver is also the organ with the highest GSH levels^81^, and, as in root regeneration, liver regeneration is also inhibited by perturbation of GSH levels via BSO treatment^82^. Thus, the metabolic environment and core signaling properties of GSH may establish some of the competence of regenerative tissue.

The regulation of G1 by GSH import and the involvement of fast divisions in pluripotency are remarkably similar facets of regeneration in plants and animals, even if the specific mechanisms have diverged. As efforts are underway in both kingdoms to improve regeneration, the mechanisms that control rapid G1 are promising tools to control the process. Our study points to a remarkably conserved role for GSH and its role in G1 truncation and highlights the role of the metabolic environment in regeneration.

### Limitations of the Study

Several corroborating lines of evidence supported our localization of GSH in the root and we used multiple methods to validate cell cycle reporters. Nonetheless, first, we point out that this work relies on dyes to visualize GSH in vivo rather than direct visualization. While direct visualization of GSH is possible via mass spectroscopy imaging, the spatial resolution of this technique is not yet fine enough to achieve cell type-specific resolution in the Arabidopsis root, where many cells are smaller than 10 microns in the x and y dimensions. Further, direct GSH biosensors are not currently available for plants. It will be important to examine GSH localization directly via live imaging when the requisite technology becomes available. In addition, an inducible inhibition of GSH production in the ground tissue would further increase confidence that the ground tissue is the source of GSH to facilitate G1 exit during regeneration. Another limitation relates to our isolation of cells by phase using Fluorescence Activated Cell Sorting (FACS). In the ideal case, we would have used the cell cycle readout of PlaCCI using FACS to define cell cycle phase to obtain bulk protoplast populations using the markers from each phase alone from the same batch of roots. However, we found that the CDT1a and CYCB1;1 fluorescent fusion proteins that mark G1 and G2/M phases in the PlaCCI reporter rapidly diminished in protoplasts. In addition, we could not directly alter G1 duration independently of other mechanisms. The direct manipulation of G1 duration would further show a role for fast G1s in regeneration. When such tools become available, they will be a valuable addition to this literature. Finally, the work does not address how rapid vs. slower reprogramming could provide an advantage to the plant. Further work could focus on the ecological or physiological advantages or tradeoffs of rapid cellular reprogramming in regeneration.

## Acknowledgements

This work was supported by a Postdoctoral Research Fellowship in Biology from the National Science Foundation (2109634) to L.L. and a National Institutes of Health grant (R35GM136362) to K.D.B. We acknowledge Dominique Bergmann, Michael Raissig, Ximena Anleu-Gil, Martin Bringmann, and Joseph Cammarata for helpful discussions.

## Author contributions

L. R. L.: Conceptualization, Methodology, Validation, Formal analysis, Investigation, Writing - Original Draft, Writing - Review & Editing, Visualization. B. G.: Resources, Data Curation, Writing - Review & Editing. R. R.: Investigation, Writing - Review & Editing, Visualization. C. H.: Investigation. B. D.: Resources, Writing - Review & Editing. C. G.: Resources, Writing - Review & Editing. K. D. B: Conceptualization, Methodology, Writing - Original Draft, Writing - Review & Editing, Supervision, Funding acquisition.

## Supplemental Figure Titles and Legends

**Figure S1.**
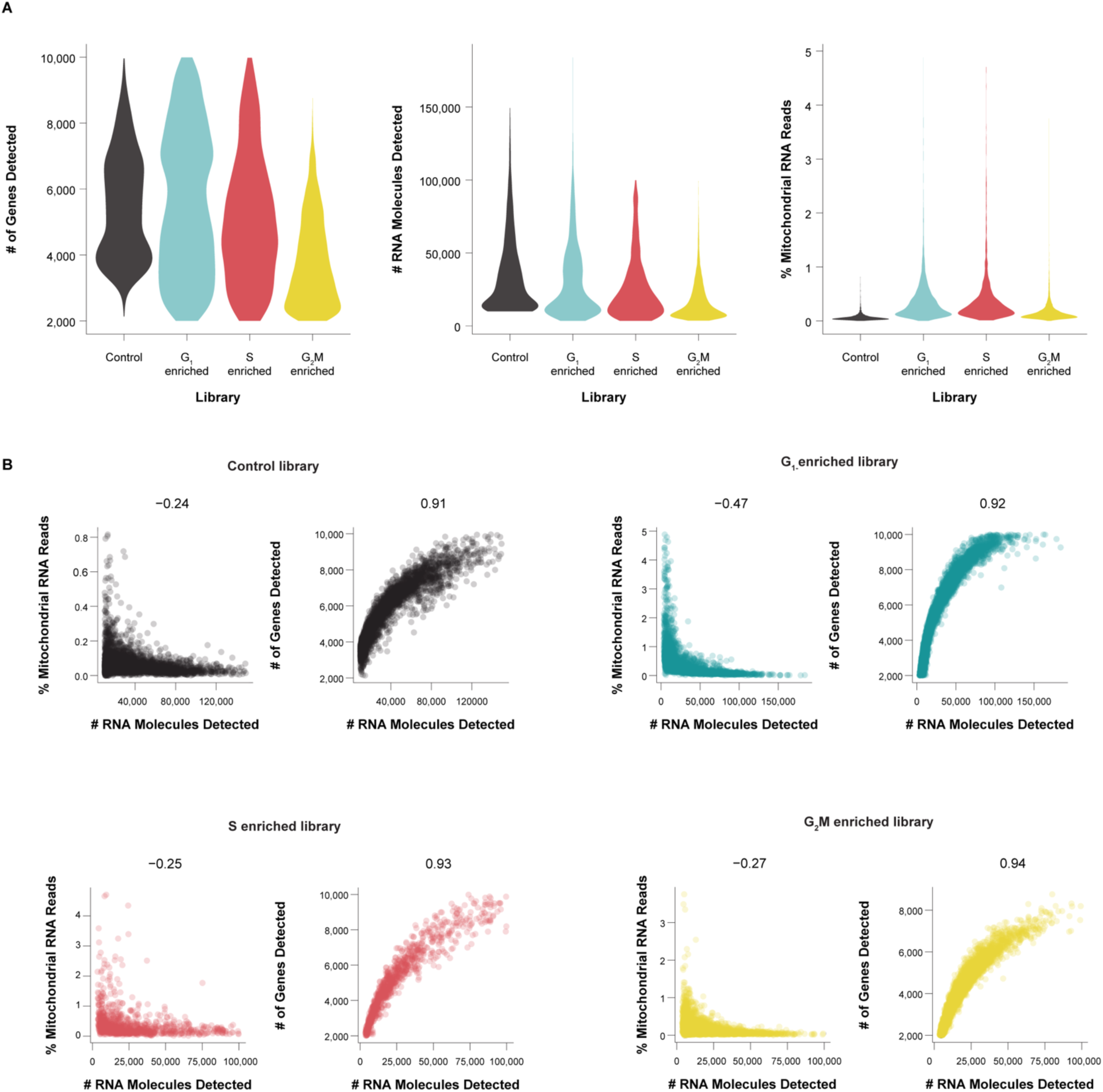
Single Cell RNA-seq Profiles Show Robust Signals in Quality Control; Related to Figure 1. (A) Violin plots showing the number of genes, RNA molecules, and the percentage of reads from mitochondrial genes, per cell in each scRNA-seq library. (B) For each library, a pair of scatter plots shows (1) the anti-correlation between percent mitochondrial reads and number of RNA molecules detected (at left), and (2) the correlation between the number of genes and the number of unique RNA molecules detected (at right). Correlation coefficient is shown above the plot.

**Figure S2:**
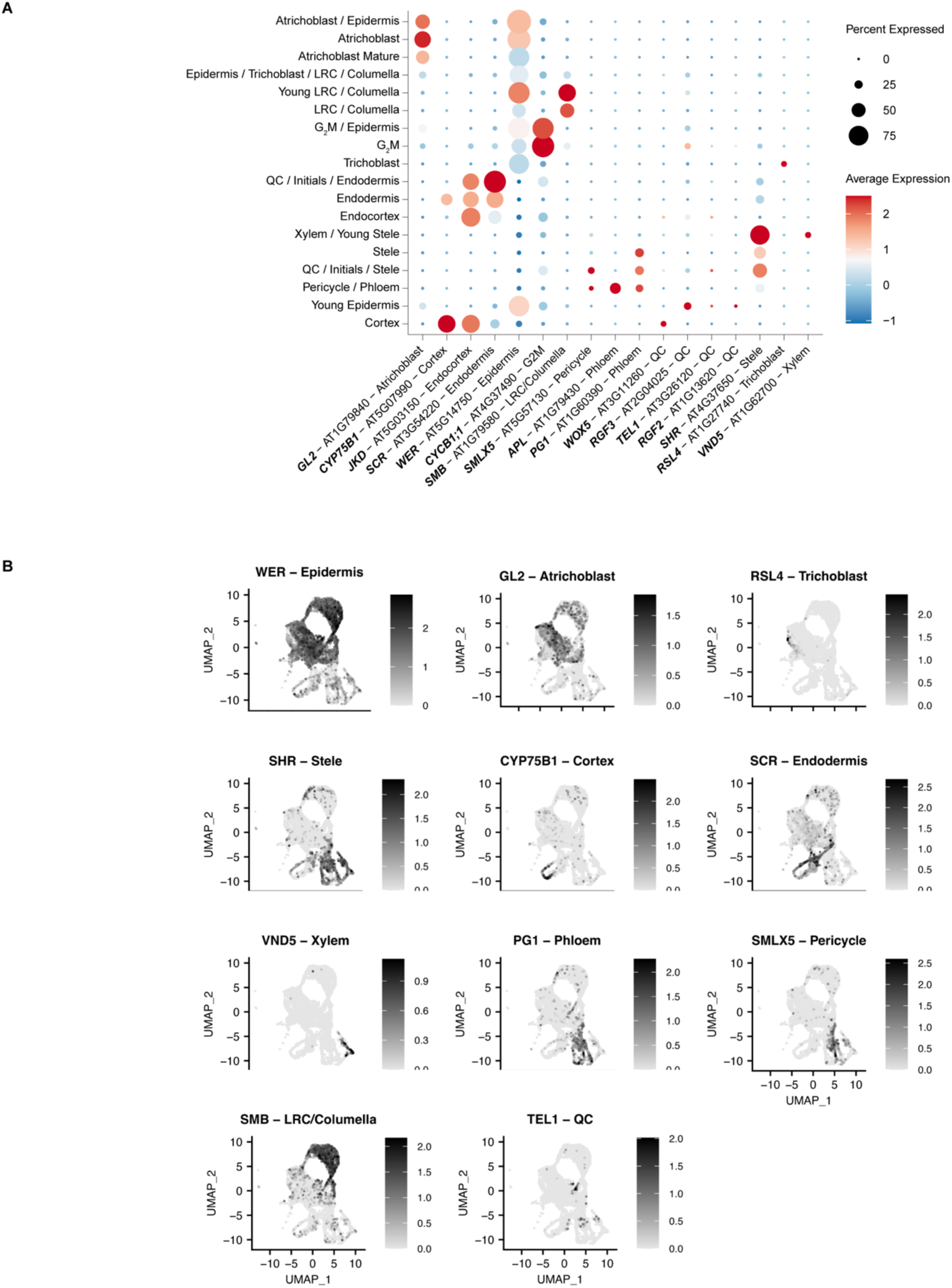
Markers robustly identify cell types in phase-enriched libraries, Related to Figure 1. (A) A dot plot showing the expression of marker genes across clusters defined by cell type in the integrated phase-enriched libraries. Size of the dot shows the percentage of cells in a cluster expressing the marker and the colormap shows the average expression of the marker in the cluster. (B) UMAPs highlighting the highly localized expression of various cell-type specific marker genes, as expected for robust capture of cell identities in scRNA-seq profiles. The cells are not grouped by phase and demonstrate the overall quality of the cells in the ability to capture clusters with clear cell identity.

**Figure S3.**
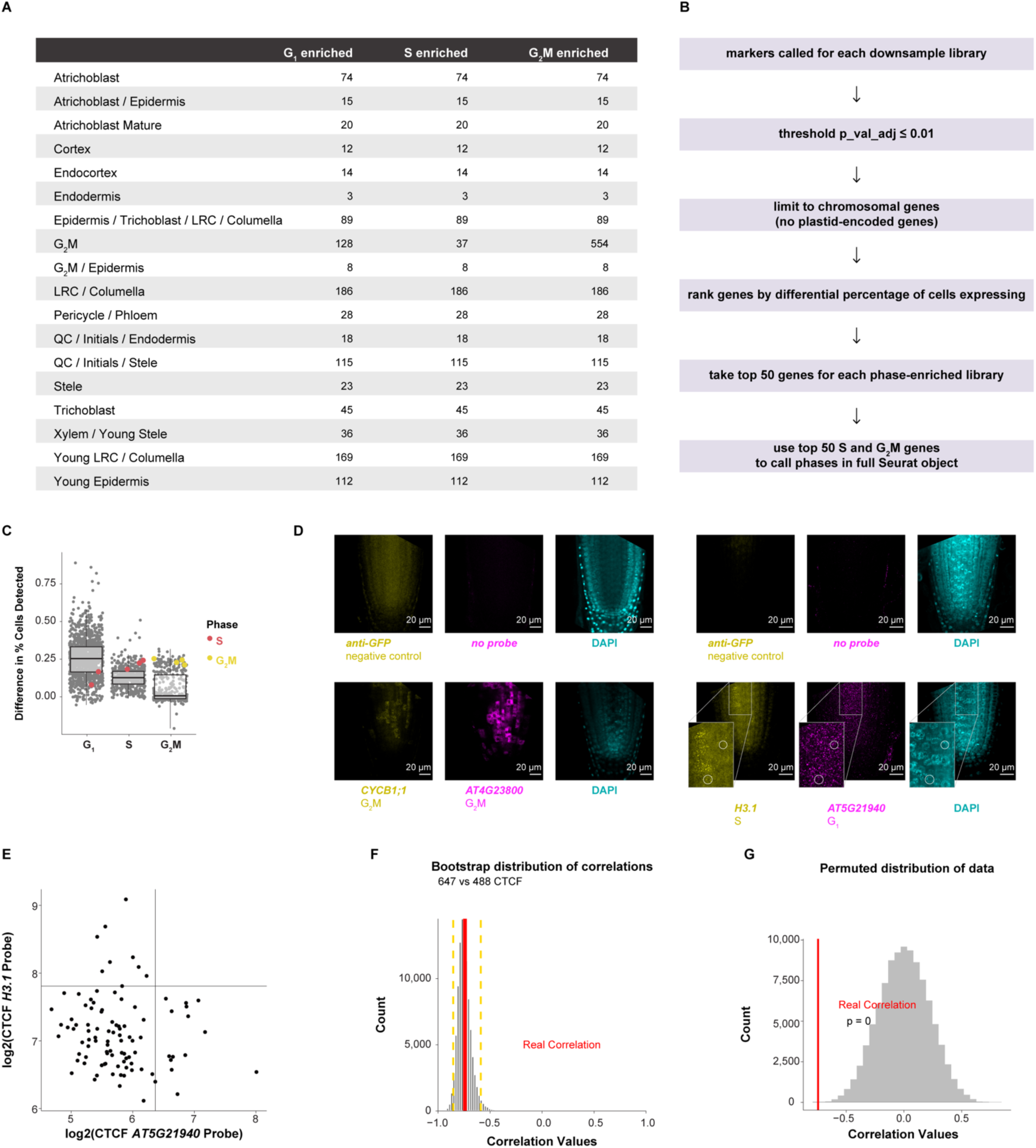
Data analysis methods identify cell phase markers with *in situ* validation of a new G1 marker; Related to Figure 1. (A) Cell counts for down-sampled phase-enriched libraries, ensuring each cell type contributed an equal number of cells to each phase enrichment analysis and each cell type contributed to phase enrichment analysis. (B) Differential expression analysis pipeline to identify phase markers. (C) Genes (each dot) categorized as differentially expressed in specific phase-synchronized libraries. The y axis represents the difference in the percentage of cells in which the gene is expressed in target versus non-target libraries. The highlighted genes are gold standard markers of phase-specific expression, showing high expression in many cells in the appropriate phase-synchronized library (x axis categories). (D) Representative images of G2M (left) and S phase (right) from *in situ* experiments shown with their corresponding negative controls as annotated. Insets highlight examples of cells where G1 and S probe signal is anti-correlated, which is quantified in the next panel. (E) Anti-correlation with signal cutoffs shown for H3.1 and AT5G21940 probes with signal cutoffs determined empirically via change point analysis^83^. Values come from three root median sections in which all cells were hand segmented based on DAPI counterstain. (F) Bootstrap distribution of correlation values between H3.1 and AT5G21940 probe signals shows the determined anti-correlation falls within the 95% confidence interval (yellow dotted lines). (G) Permutation distribution of the correlation between H3.1 and AT5G21940 probe signals shows the actual anti-correlation falls well outside of the null distribution (pvalue = 0).

**Figure S4:**
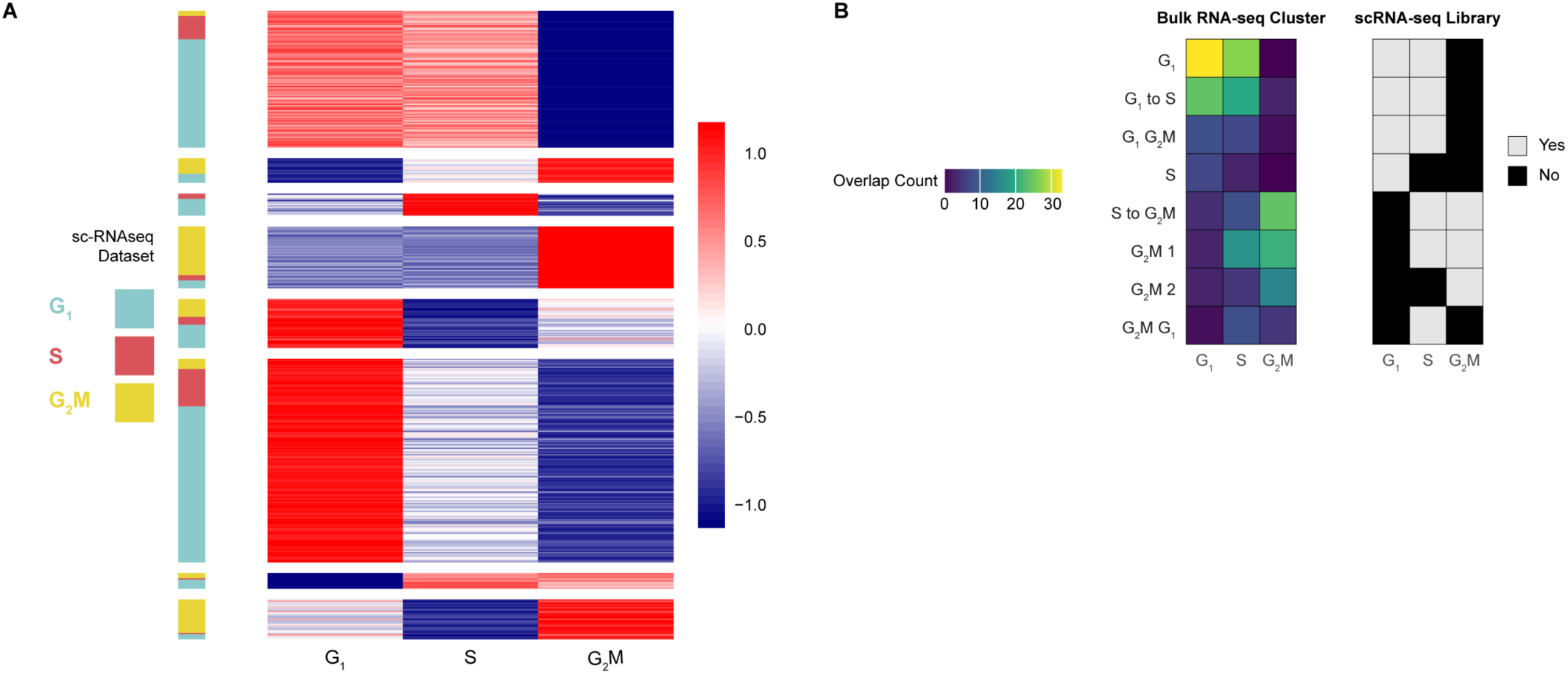
Bulk RNA-seq profiles of the cell cycle confirm phase-enriched scRNA-seq; Related to Figure 1. (A) Gene expression heatmap (red and blue) in which each row is a gene and each column represents the average expression profile across bulk RNA-seq profiles where the three libraries represent cells sorted by ploidy level as a proxy for phase. The color bar to the left indicates the independent cell cycle phase classification of each gene from analysis of the synchronized scRNA-seq library. In the bulk RNA-seq analysis, genes were grouped into 8 k-means clusters. Agreement between the two independent analyses is indicated by groups of genes showing a sc-RNA-seq classification and enrichment in the appropriate ploidy sorted cell library. Strong agreement is shown for G1 and G2/M, while S-phase is not well defined in the ploidy sorting (B) Heatmaps showing the number of overlapping genes (left) and the statistical significance of the overlap (right) between differentially expressed genes from phase-enriched scRNA-seq (columns) and gene expression clusters of ploidy-sorted cells determined by k-means clustering (rows). Yes=statistically significant overlap at p<0.05 by Fishers exact test. See also Table S4.

**Figure S5:**
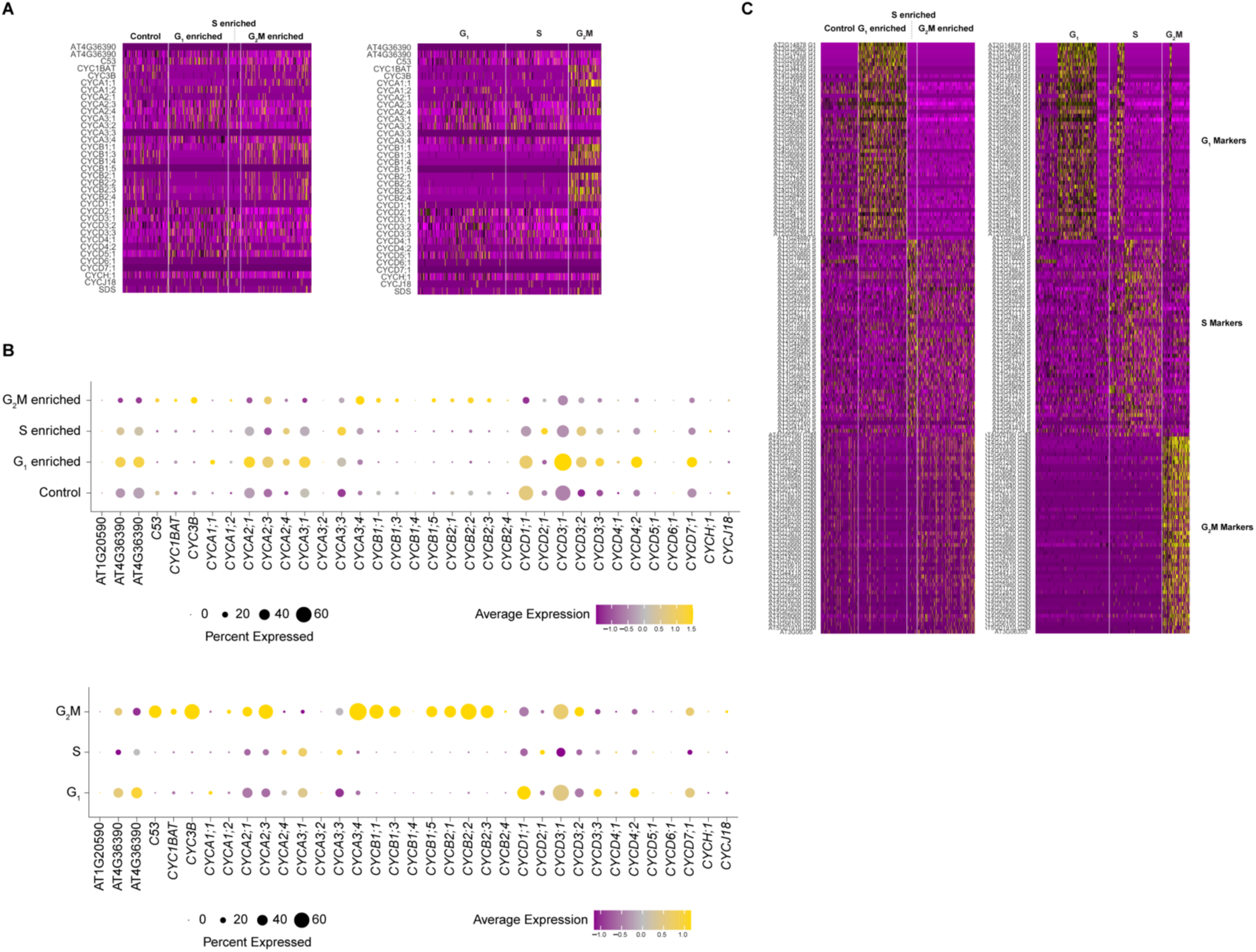
Enrichment analysis for phase markers shows agreement with known cell cycle markers but identifies more robust markers; Related to Figure 1. (A) Heatmaps comparing expression of classical cell-cycle markers (rows) in cells (columns) grouped by the phase enrichment library from which they came (left), which may still contain cells from a mixture of phases, vs. cells assigned to phase based on marker analysis (right). At left, some enrichment of markers is visible but phase enriched libraries still contain cells in the non-target phase. At right, enrichment of known markers is more prominent when cells are assigned to phase by our analysis pipeline, which is independent of the expression of the classical cell cycle markers. (B) A summary analysis of the heatmap data in A. Dotplots show the expression of cyclins in phase-enriched libraries (top) vs phases assigned with our top marker genes (bottom). Cyclins are expressed in the appropriate datasets despite their sparseness (top). Cyclin expression behaves well based on phase assignments performed with our marker genes (bottom). (C) Following the same comparison as in A with the top 50 markers assigned by our pipeline. At left, the markers are shown based on their enrichments in the different phase libraries. These agree with classical markers but the analysis shows the new markers have higher expression and are more frequently detected in single-cell profiles. At right, the analysis show cells classified by phase using the top 50 markers. Note that many G1-phase markers also express in early S phase, but S-phase has distinct markers to separate G1 and early S.

**Figure S6.**
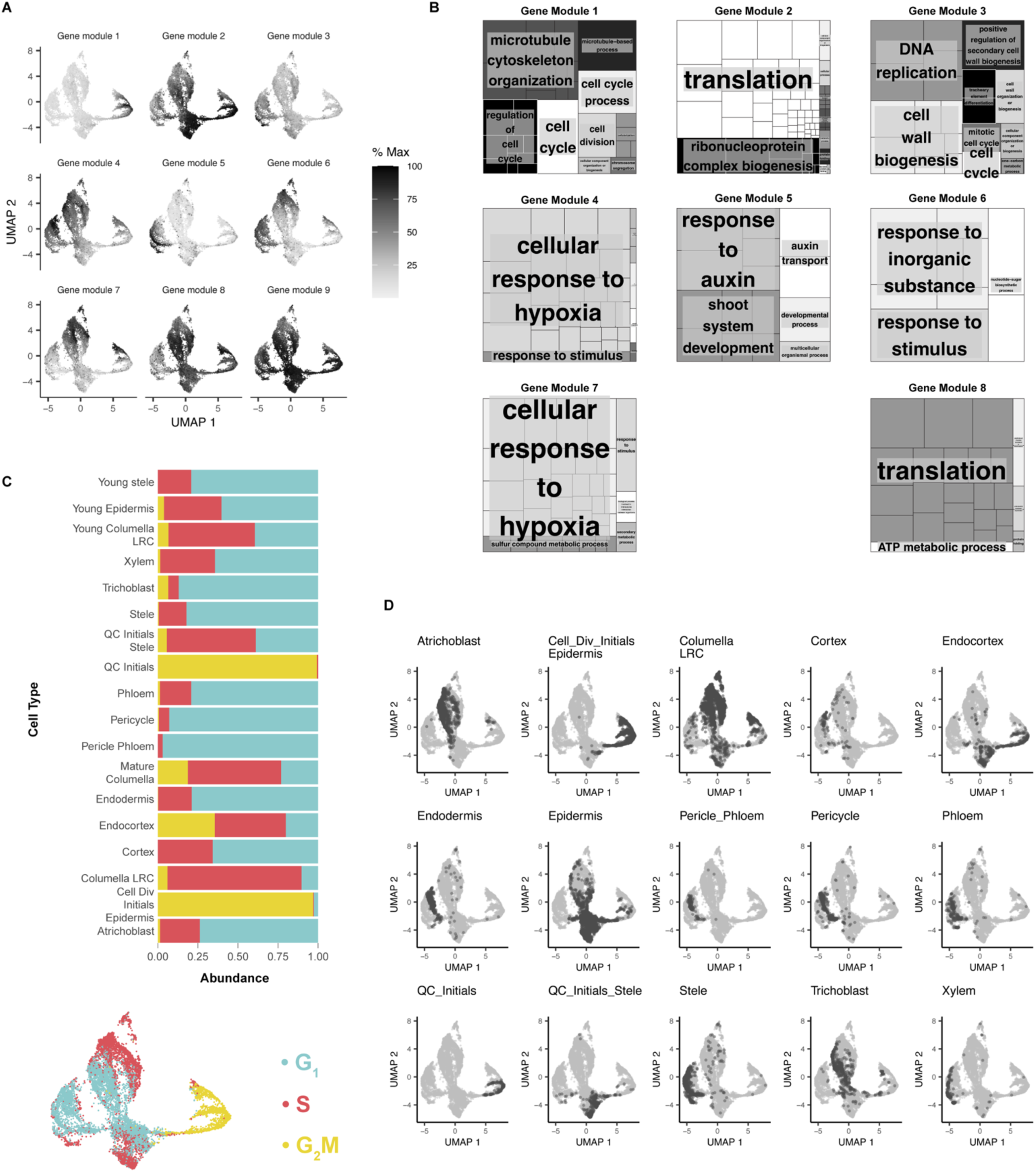
Cells of the same identity group together even when clustered by only cell cycle markers; Related to Figure 2. (A) Analysis of gene modules that are preferentially expressed along the cell-cycle pseudotime ordering, as determined by Monocle3 (see Methods). Grayscale shows the aggregate gene expression of each gene group. (B) GO-terms associated with the corresponding gene group shown in B. No significant GO terms were found for gene module 8. (C) Relative abundances of phases among each cell type are shown. (D) UMAP outputs of pseudotime analysis clustered using the top 50 cell-cycle markers with an independent analysis of cell identity mapped onto the UMAP trajectories. In each panel, a different cell type is highlighted in red. At left, a key shows the cell cycle classification for each cluster in the UMAP.

**Figure S7:**
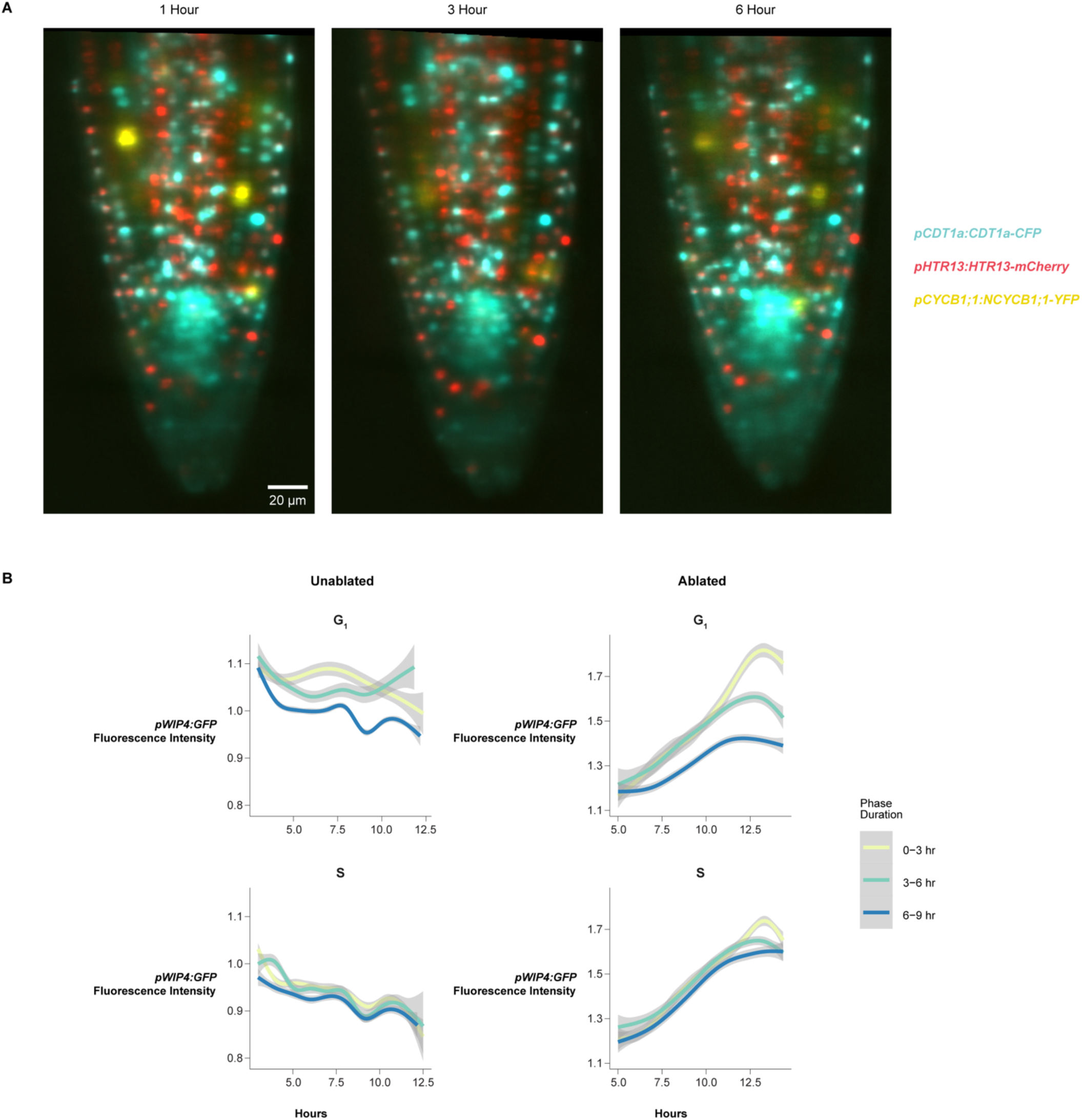
The appearance of newly reprogrammed cell identity correlates with rapid G1 phases; Related to Figure 3. (A) Representative images of a control root expressing PlaCCI and *pWIP4:GFP* at 1, 3, and 6 hr time points during a time-lapse acquisition, showing consistent distribution of each of the three markers over time under imaging conditions (B) Quantification of the WIP4 signal intensity in G1 phase and S phase cells over the duration of time-lapse movies. The figure represents the complete analysis of data shown in Figure 3E.

**Figure S8.**
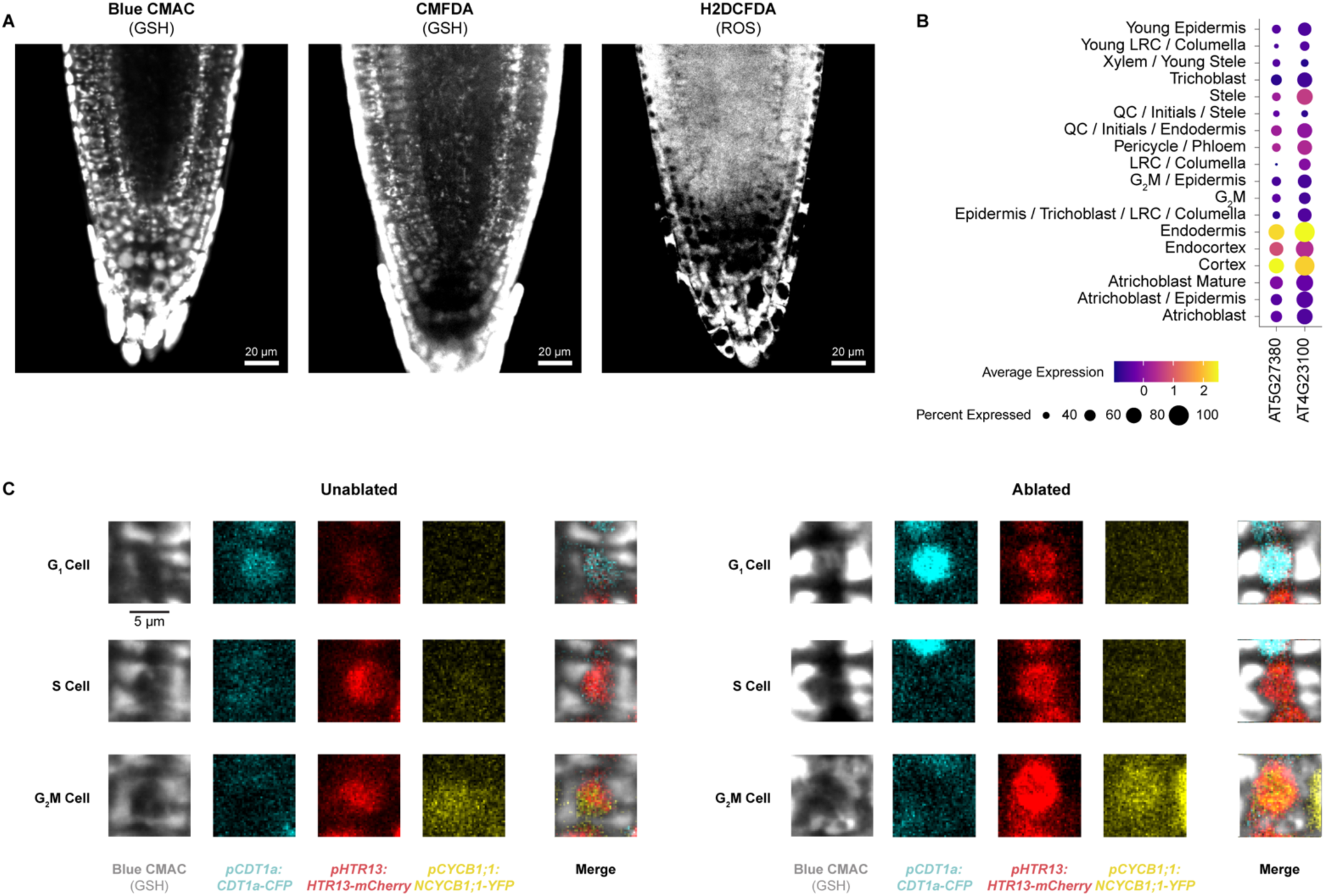
ROS and GSH dyes show different tissue localization patterns; Related to Figure 4. (A) Representative confocal microscopy images of seedlings stained for GSH (Blue CMAC, CMFDA) or ROS (H2DCFDA) under control conditions. Note that the two GSH dyes agree and show prominent ground tissue staining. Note that CMFDA and H2DCFDA, with similar chemical structure but different target molecules, show different staining patterns. (B) Expression of GSH1 and GSH2 represented as dot plot derived from scRNA-seq profiles in different root cell types. Note the prominent expression in endodermis and cortex, in agreement with the GSH dyes. (C) PlaCCI signal at time 0 for cells shown in Figure 4D, which demonstrates a GSH burst in G1 nuclei prior to fast divisions. Exogenous application of GSH did not cause a shift in the number of G1 cells (root n = 37, nuclei n = 9100, no significant difference between treatment and control by student’s t test).

**Figure S9.**
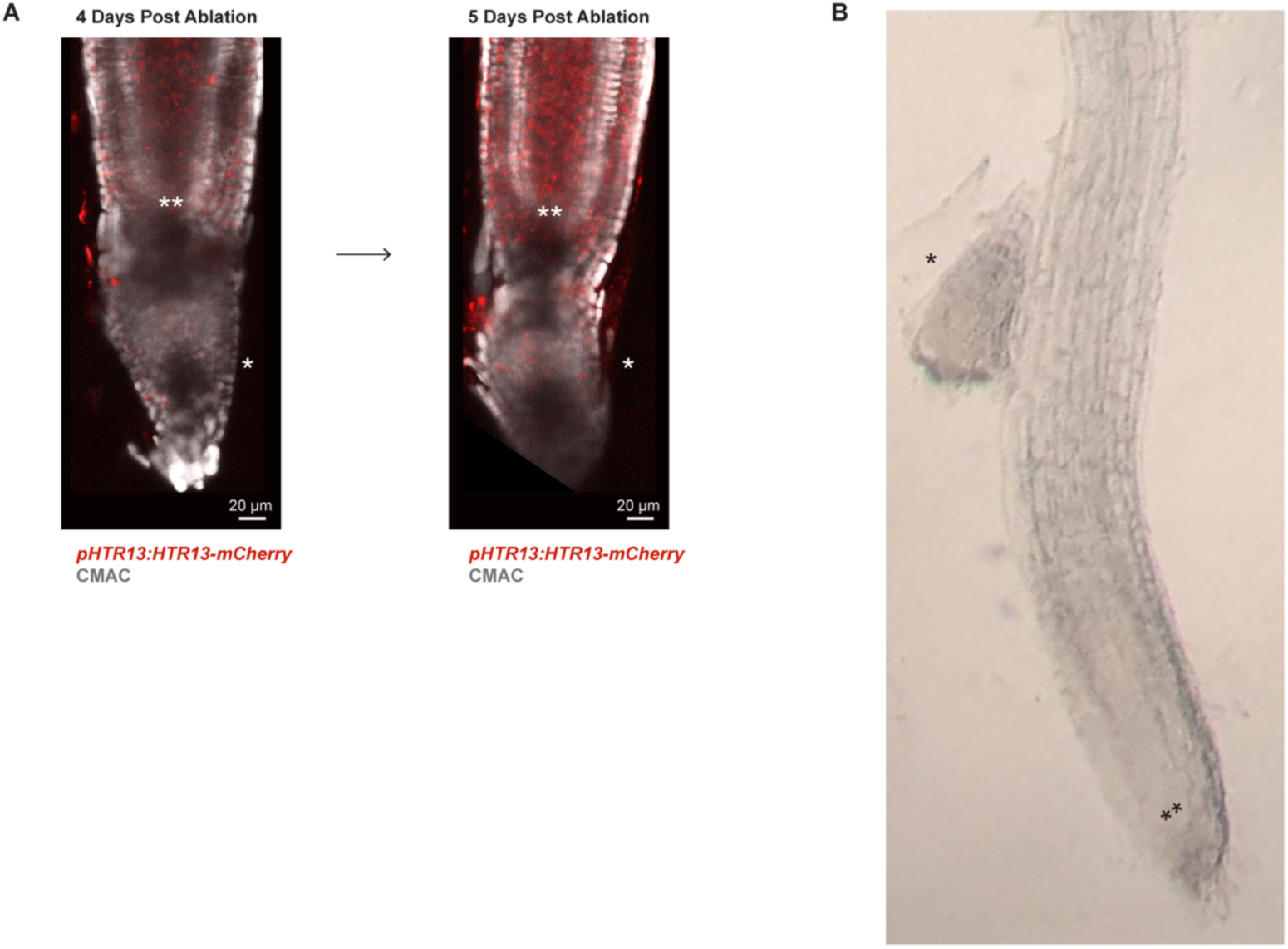
Transverse ablation leads to the reformation of a new root tip similar to the root tip excision procedure; Related to Figure 4. (A) Representative confocal images of seedlings (grown on standard ½ MS and then mounted in an imaging cuvette) undergoing regeneration. Between days 4 and 5 post ablation it becomes apparent that new columella above the ablation is established proximal (shootward) to the original QC (*), which is below the ablation. The tapered root cap, which includes the columella, is apparent distal to the new QC (**), both of which are above the ablation site. (B) At a later time point, the original root tip (*) is sloughed off as growth continues from the new QC/stem cell niche (**) in the same seedling shown in the lower panel of A.

**Figure S10.**
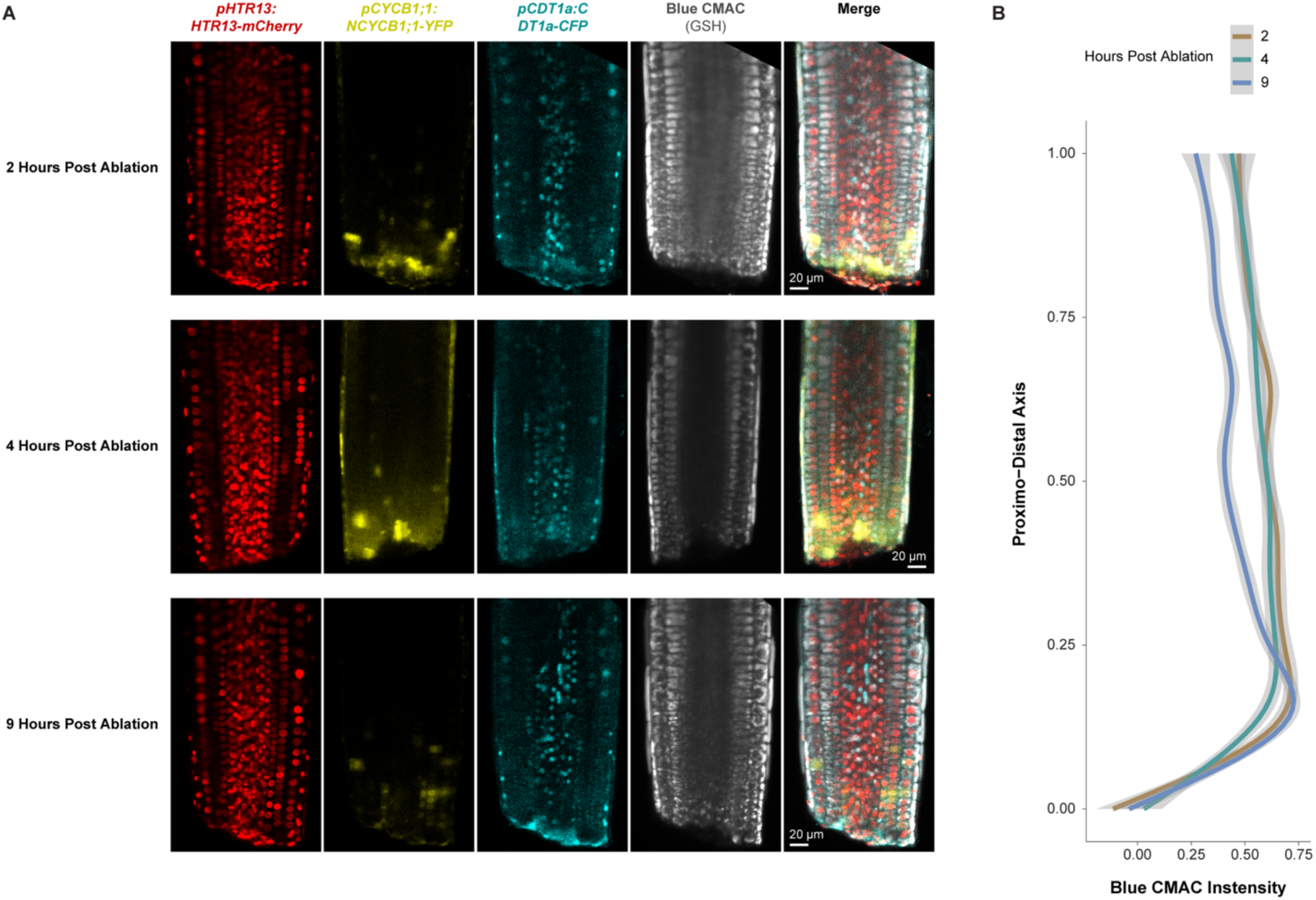
GSH dye CMAC is brightest in the same region where cells undergo rapid division and shortened G1 during regeneration; related to Figure 4. (A) Representative confocal images of PlaCCI roots stained with Blue CMAC. Images were taken 2, 4, and 9 hpc. (B) Quantification of nuclear CMAC staining intensity along the proximal-distal axis at different time points after ablation. The y-intercept represents the ablation site and the range of the y-axis represents the visible length of root imaged in the frame as shown in A. Note the peak of CMAC intensity right above the cut site between 0.00 and 0.25 on the longitudinal axis of the root (y-axis), which is highest at 2-4 hr post cut and begins to dissipate above point 0.25 at 9 hr.

## STAR Methods

### Resource Availability

#### Lead contact

Requests for resources or plant lines should be addressed to the lead contact for this work, Kenneth Birnbaum.

#### Materials availability

Arabidopsis lines generated for this work are available following publication upon request.

### Experimental Models and Study Participants

#### Plant growth and treatment conditions

Arabidopsis col-0 seedlings were grown vertically in an incubator set to long day conditions on ½ MS media unless otherwise noted. For hydroxyurea (HU) treatment, seedlings were synchronized in one of three cell cycle phases as previously described^36^. Briefly, seedlings were grown until 6 DPG vertically on ½ MS on top of sterile mesh (product #03100/32, ELKO Filtering Systems). Then seedlings were transferred to MS plates supplemented with 2mM HU (product # H8627, Millipore Sigma). Various incubation times were used to synchronize cells in different phases of the cell cycle as follows: 6 hr for S phase, 17 hr for G2/M, and 22 hr for G1. Synchronization in each phase was confirmed via confocal microscopy using the PlaCCI reporter. For BSO treatment, seedlings were germinated on MS media alone (control) or supplemented with 1 or 0.5 mM BSO (product # B2515, Millipore Sigman) as previously described^31^. Seedlings were grown vertically on this media until they were 7 DPG and then used for either imaging or regeneration assays. Regeneration assays were performed by manually removing the distal-most 70 microns of the root tip using an ophthalmic scalpel (product #72045-15, Feather Safety Razor Company). Roots were then allowed to grow while regeneration was monitored by either staining for amyloplasts at 18 hr with mPS-PI^50^ or by counting the proportion of roots that had recovered gravitropism at 48 hr^2^. PlaCCI seedlings (*pCDT1a:CDT1a-eCFP*, *pHTR13:HTR13-mCherry* and *pCYCB1;1:NCYCB1;1-YFP*, where “N” denotes an N terminal fusion) were crossed to cell type reporters including *pWIP4:GFP* (columella and QC), *pWOX5:YFP* (QC), *PET111:YFP* (mature columella), and *pSCR:erYFP* (endodermis and QC). An estradiol inducible callose synthase line^54^ driving induction in the cortex and endodermis was used for plasmodesmatal block experiments. For these experiments, plants were grown vertically on sterile mesh on top of ½ MS for 7 days, then transferred to ½ MS supplemented with 1 µM estradiol for 17 hr. Where noted, estradiol plates also included GSH (0.5mM) or sucrose (1%). Plants were then transferred back to unsupplemented ½ MS. Regeneration experiments were then performed as described above. For root growth experiments, root tip locations were marked after transfer back to ½ MS and then growth from that point was measured 24 hr later.

All experiments, unless otherwise noted, were performed on seedlings at 7 dpg.

### Method Details

#### Confocal microscopy

Multichannel imaging was performed on a Zeiss 880 Airyscan microscope. Channel acquisition parameters were initially defined using the Zen Smart Setup feature and then refined to ensure the acquisition range was narrow and centered over the emission peak. Channels were then acquired in sequential scans to maximize signal and minimize spectral overlap.

Laser ablations that were sufficient to cause new meristem establishment (regeneration) were performed using a Coherent Chameleon Vision II 2-photon laser on a Zeiss 880 Airyscan microscope. A 2-dimensional ROI was specified using the Zeiss ROI manager in the Zen Acquisition Black software with the time series, bleaching, and ROI modes enabled. This ROI targeted a transverse section of the root that was positioned approximately 10-20 microns shootward of the QC that spanned the entire medio-lateral dimension of the root with a thickness of approximately 5-10 microns. The ablation laser was used at 710 nm at 100 percent power for 15 iterations. In order to ensure sufficient tissue damage was achieved to induce the root to establish a new meristem, the ablation was performed in 3 Z planes: (1) in the medial plane, and then on both sides of the medial plane (2) closer to the cover slip and targeting the epidermis and cortex (about 15-20 microns off the medial plane), and (3) further from the cover slip than the median plane as deep as the confocal microscope could image into the tissue before imaging quality degraded (15-20 microns from the medial plane). Each ablation was performed as part of a time lapse acquisition, in which typically two frames were acquired, followed by the ablation, and then three additional frames were acquired. These frames were set to be acquired 1 millisecond apart, which functionally resulted in continuous acquisitions and total time lapses of approximately 90 seconds. For 30-minute-long time lapses taken on the Zeiss 880 Airyscan confocal, frames were acquired in one Z plane three minutes apart. This laser ablation strategy was adopted to enable imaging of injured roots that were already mounted in a cuvette compatible with our light sheet setup (described below) so that we could monitor injury response via time lapse microscopy without any confounding effects of the stress of mounting seedlings after root tip removal.

Plants were stained with Blue CMAC by mounting in imaging cuvettes as described above using media supplemented with Blue CMAC (ThermoFisher #C2110) to achieve a concentration of 10 µM once the media had equilibrated to 30 degrees Celsius. Media was then split into a number of batches equal to the number of treatment conditions to ensure that all conditions received the same concentration of Blue CMAC. Additional treatments were then supplemented into the relevant batch of media as required. 5 mL of each media treatment was then added to its own cuvette and cured for at least four hr at 4 degrees Celsius. Plants were then transferred to an imaging cuvette and allowed to recover in the growth chamber overnight. For CMFDA and H2DCFDA staining, seedlings were transferred to liquid ½ MS supplemented with either stain to a final concentration of 1 µM for 1 hr prior to imaging.

#### Light Sheet Microscopy

All time lapse movies were performed on an inverted Leica model Dmi8 outfitted with a Tilt Light Sheet Imaging System (Mizar) with filters optimized to visualization of YFP, CFP, and mCherry (Chroma). All roots were imaged at 7 dpg. Samples were mounted for light sheet microscopy as follows: plants were grown vertically on MS plates for 6 days. On day 6, 5 mL of MS with 2% low melt agarose was cast into imaging cuvettes (CellVis product number #C1-1.5H-N) after being filtered through a 0.45 micron nylon filter (product # 76479-042, VWR) to remove any particulates that might disturb the path of the light sheet to prepare media “blankets”. These blankets were stored at 4 degrees Celsius for at least four hr prior to mounting to ensure they had fully polymerized. A sterile scalpel and forceps were used to remove a small amount of media from one end of the cuvette to create a gap that could be used to lift the media out of the cuvette. The scalpel was then gently run along the edge of the imaging chamber to free the blanket while producing minimal distortions to the media. Sterile canted forceps were then used to gently lift the media blanket out of the cuvette and placed in a sterile petri dish. Several 6 DPG seedlings were placed on top of the media blanket such that the roots were in contact with the blanket and the shoots hung off the edge. A fresh cuvette was then lowered over the blanket until the blanket made contact with the cover slip at the bottom of the cuvette. Seedlings were inspected for tissue damage under a brightfield microscope and any gaps between the blanket and the wall of the cuvette were filled in with additional filtered media prepared as above to ensure the light sheet did not pass through any air gaps. The assembled cuvettes were then placed into a growth chamber overnight oriented such that the roots pointed downward to allow the plants to recover from the stress of the mounting procedure. Roots were imaged with a 40X water immersion objective, with stacks spanning the entire Z dimension spaced 1.5 microns apart acquired every ten minutes in mCherry, CFP, and YFP to create time lapse movies of PlaCCI. Laser power and acquisition time was adjusted for each experiment to account for variable distance of the sample to the side of the cuvette through which the light sheet enters. A sample binning of 2 was used to improve signal brightness. For imaging of the F3 progeny of PlaCCI crossed to the WIP4 transcriptional reporter or the *PET111:YFP* enhancer trap line^48^ in which both transgenes had been screened for stable brightness, a fourth channel - GFP - was imaged. No photobleaching was observed using these imaging conditions over the course of a time lapse. To maintain imaging quality, water was added to the 40X objective after 7-10 hr of imaging depending on the ambient humidity. This was accomplished by briefly removing the imaging cuvette between acquisitions, adding additional water to the objective, and then replacing the cuvette. The stage was adjusted to recenter the sample and then the image was realigned *post hoc* using Imaris to account for any subtle shifts in sample position. This allowed us to avoid moving the stage, which would necessitate adjusting the focus of the light sheet midway through the time lapse acquisition.

#### scRNA-seq

Protoplasts were generated as follows: To collect roots enriched for different phases of the cell cycle, root tips were synchronized with 2mM HU media as described above. To process cells synchronized in different phases in parallel, seedlings were transferred to HU media in a staggered manner such that they would be ready for harvesting at the same time.

The distal-most 400 µm of approximately 500 root tips were excised from 7 DPG seedlings and then collected via capillary action with a P200 pipette tip containing 25 µL of protoplasting buffer. These root tips were then dispensed into cell wall degrading solution as previously described^84^. Root tips were gently agitated on an orbital shaker for approximately 1 hr and were gently pipetted up and down with a P1000 pipette every ten minutes after the first half hr of incubation. Root tips were then passed through a 40-micron cell strainer (product # 08-771-1, Fischer Scientific) and any large aggregates of cells were gently pressed against the strainer using sterile flat forceps to release any cells that had so far failed to dissociate.

10X libraries were prepared from protoplasts to generate scRNA-seq libraries using the Chromium Next GEM Single Cell 3ʹ Reagent Kit v3.1 (10X Genomics) following manufacturer’s instructions.

The cDNA and sequencing library fragment sizes were both measured with the Agilent Tapestation 4200 using the high sensitivity 1000 (product # 5067-5582, 5067-5583) and 5000 (product # 5067-5592, 5067-5593) reagents respectively. Sample concentration was detected using the Qubit HS dsDNA (product # Q32851, Thermofischer) assay following manufacturer’s instructions. Library quantitation for pooling was performed as follows: the fragment size and concentration of the library in ng/µL were used to determine the molarity of the libraries with the following equation: [Lib Conc (ng/µL)]/[(Frag Length (bp) * 607.4)+157.9] * 1000000. Libraries were then diluted to 3 nM concentration and pooled for sequencing. Samples were sequenced on a Novaseq 6000 using an SP flowcell in 28x91 paired end 100 cycle mode with V1.5 reagents (100 cycles).

#### Bulk RNA-seq

For bulk RNA-seq, total RNA was extracted from sorted protoplasts using the Qiagen RNA micro kit following manufacturer’s instructions. RNA quality was determined using RNA high sensitivity reagents (product # 5067-5579, 5067-5580, 5067-5581, Agilent) for the Agilent TapeStation 4200. Total RNA was used to synthesize cDNA using the SMART-Seq v4 full-length transcriptome analysis kit from Takara (product # 634888) using protocol B specified in the manual on page 12. The quality of cDNA was then assessed using D1000 reagents for the Agilent Tapestation. The resulting cDNA was used to generate sequencing libraries with the Ovation Ultralow Library System V2 from Tecan (product # 0344) following manufacturer’s instructions. Libraries were then sequenced on a Novaseq 6000 with an SP flowcell in 1x100 single end 100 cycle mode with V1.5 reagents (100 cycles).

Cells were collected by FACS as follows: Root protoplasts were sorted using a BD FACS Aria II using FACS Diva software as described previously^85,86^. Briefly, protoplasts were sorted directly from cell-wall degrading solution into a 1.5 mL microcentrifuge tube containing 350 µL of Qiagen RNA extraction buffer supplemented with beta mercaptoethanol.

Protoplasts expressing an H2B RFP fusion and a CDT1a GFP fusion under the native promoter were sorted and gated to remove doublets and debris. Then RFP positive events were identified by plotting red scale autofluorescence versus RFP and then gating for cells that showed RFP fluorescence above background as defined by a Col-0 control expressing no fluorescent proteins. In tandem, CDT1a positive cells were identified by plotting autofluorescence versus GFP and gated for GFP expression above background relative to Col-0 control. Then both the RFP+ and GFP+ populations were plotted in a histogram of RFP signal v. cell count. This revealed a population with two RFP peaks characteristic of DNA staining in dividing cells. The GFP+ population (CDT1a reporter fluorescence) overlapped with the 2n ploidy peak, which is consistent with its expression in the G1 phase of the cell cycle and was used as a positive control. Further gates were defined based on the histogram to collect cells in G1 (2n), G2/M (4n), and S (intermediate RFP signal) phases. These populations were collected simultaneously in a three-way sort and the maximum number of cells were collected for each phase. This protocol was repeated independently twice to generate 6 samples for RNA-seq library preparation. Samples were snap frozen and stored at -80 degrees Celsius until all samples were collected and could be processed for RNA extraction and library preparation simultaneously.

In order to use cellular ploidy as a proxy for cell cycle phase, it was critical to harvest the distal-most portion of the root tip in order to avoid harvesting any cells that had already begun endoreduplication. The distal-most 200 µm of approximately 500 root tips were excised from 7 DPG seedlings and then collected via capillary action with a P200 pipette tip containing 25 µL of cell-wall degrading solution. These root tips were then dispensed into cell-wall degrading solution. Root tips were gently agitated on an orbital shaker for approximately 1 hr and were gently pipetted up and down with a P1000 pipette every ten minutes after the first half hr of incubation. Root tips were then passed through a 40-micron cell strainer and any large aggregates of cells were gently pressed against the strainer using sterile flat forceps to release any cells that had so far failed to dissociate. The resulting protoplasts were then transferred to a test tube appropriate for the cell sorter and immediately processed via FACS.

#### Sequencing Data Analysis

##### Bulk RNA-seq

For Bulk RNA-seq, reads were trimmed using Trimmomatic version 0.39 in single end mode with the following settings: ILLUMINACLIP:TruSeq3-SE:2:30:10 LEADING:3 TRAILING:3 SLIDINGWINDOW:4:15 MINLEN:36. Trimmed reads were mapped to the Arabidopsis TAIR10 genome using HISAT2 version 2.2.1. Reads mapping to genes were counted with Rsubread (version 1.22.1) featureCounts in single end mode with a minimum overlap of 5 and counting only primary alignments and ignoring duplicates. Reads were normalized using the TPM calculation and the resulting count matrix was used to calculate mean values per condition, filtered to remove genes with low expression and low variance, and then clustered via k-means clustering. The number of k (8) was chosen to reflect the total permutations of expression changes (up or down) and cell cycle phases (G1, S, G2/M).

##### scRNA-seq

For scRNA-seq the mkfastq function in Cell Ranger 5.0.1 was used to generate fastq files from the raw sequencing output. Count matrices for scRNA-seq experiments were then generated with the count function and the TAIR 10.38 release of the Arabidopsis genome.

##### Quality Control – scRNA-seq

After generating count matrices using Cell Ranger, Seurat was used to filter cells based on the number of features detected (more than 2000 and less than 10000), percent mitochondrial reads (less than 5), and total RNA molecules detected (less than 100000). This produced datasets in which the R squared coefficient between features and counts exceeded 0.93, indicating that the remaining cells in the dataset were healthy singlets. Libraries were integrated using the sctransform workflow in Seurat^37^.

##### Identifying Cell Cycle Markers

Cell type annotations were carried over from a control dataset that had previously been annotated based on the expression of cell type specific marker genes. Cell labels were carried over manually by examining the cluster membership of cells from the control library, which formed the same stable clusters as they had in previously when integrated with this dataset. Previous cluster identity was then manually transferred to all cells from the HU-treated datasets that shared cluster membership with the annotated cells from the control dataset.

Transcriptional detection of phase enrichments for scRNA-seq libraries were validated by comparing upregulated genes in each scRNA-seq library with expression patterns in ploidy-sorted bulk RNA-seq. Due to the absence of a clear peak for S phase, we collected many fewer cells from S-phase. Thus, we did not expect a high overlap in this phase. However, phase agreements were high in both G2/M and G1 phases, validating the synchronization method. For S phase, upregulated genes in the enriched scRNA-seq libraries were enriched for functions already known to be core for S-phase including many histones. Thus, we used the scRNA-seq to generate markers because of its high resolution of each phase.

While the scRNA-seq libraries were enriched for cells in each phase of the cell cycle, their cell type composition was variable. To ensure the identification of cell cycle markers present in all cell types, we projected them on the same UMAP space, determined the lowest number of each cell type across all enriched libraries and then randomly down sampled each cell type in each library to produce libraries with equal cell type composition. We then performed differential expression analysis with cells from each phase enriched library using Seurat’s FindAllMarkers function. Markers were ranked by percent differential expression and the top 50 for each library were chosen as cell cycle marker genes. Markers were then used to analyze the cell cycle in the full (not down sampled) scRNA-seq dataset and other non-synchronized scRNA-seq datasets.

In a separate analysis, we isolated individual cell types from the scRNA-seq dataset, grouped cells by phase and then performed a differential expression analysis to identify phase markers on a per cell basis. We filtered out genes with a p value great than 0.000001 and then constructed a cell type+phase by gene matrix, where each cell of the matrix contains a 1 if a gene is a marker for that cell type+phase combination, or a 0 if it is not a marker. That matrix is provided as Table S6.

##### Pseudotime Analysis

For cell cycle psuedotime analysis, Monocle3 was used to create the UMAP embeddings with the top 150 ranked genes for each phase of the cell cycle. We then used the learn_graph and order_cells functions to calculate a pseudotime trajectory for cells based on the cell cycle. To find genes that changed as a function of pseudotime we used the graph_test function. We then aggregated the gene expression matrix based on evenly spaced bins along the pseudotime trajectory and clustered those bins based on gene expression to assign genes to different positions in the pseudotime trajectory.

Data visualization was generated using ggplot2 with Tidyverse, Seurat, pHeatmap, Treemap and Monocle3.

#### Imaging Data Analysis

Long-term time-lapse images were registered in 3 dimensions by first detecting objects (either nuclei, WOX5, or WIP4 marker expression) and then using detected objects to correct the reference frame for the time lapse in 3 dimensions. The new reference frame was then used to correct the time lapse for both translational and rotational drift. Once drift corrected, nuclei were then segmented again using the spot detection tool with the local contrast setting enabled to account for uneven background throughout the root or bleed through from other channels. Once segmented, statistics for all nuclei were exported to R for further analysis. Cell phase was determined by measuring the amount of YFP, CFP and mCherry signal in each nucleus. If CFP or YFP signal exceeded a detection threshold cutoff, cells were classified as G1 or G2M respectively. All other cells were classified as S phase. The PlaCCI reporter does not easily distinguish between cells in S phase versus early G2, so it is possible that some G2 cells were classified as S phase cells in this analysis. Counts of cells in G1 (Figure 3B), G1 durations (Figure 3D), and G1 exit time (Figure 6B) were determined manually. The log rank test was used to determine the significance of the G1 survivorship analysis^87,88^.

For still images, 3-dimensional segmentation was performed in TrackMate by treating the Z dimension as a time dimension. Nuclei were segmented based on the HTR13-mCherry channel and then data for each channel within nuclei was exported to R for further analysis. In the case where a single slice was taken, all nuclei were similarly segmented in TrackMate in one dimension and retained.

Confocal image stacks were taken such that nuclei would appear in at least two consecutive slices. Therefore, all nuclei that appeared in only one slice were discarded. For the remaining nuclei, or for all nuclei in the case of images acquired as median slices, Blue CMAC signal was scaled from 0 to 1 per cell file to render nuclei comparable. In the case of short-term time lapses of PlaCCI roots stained with Blue CMAC taken using confocal microscopy, drift was corrected in 2 dimensions using the Correct 3D drift plugin in FIJI prior to Trackmate segmentation. Nuclei were filtered if they were not tracked for the entire time lapse. Blue CMAC signal was calculated as a change over the value at time zero. Tissues were classified into specific identities for quantification of GSH content using relative cell position and root morphology.

#### In Situ Hybridization

Probe selection - Candidate probes were selected from the top marker set described above if they had a were expressed in at least 80 percent of cells from the target phase and if they exceeded a differential expression threshold of 0.25 LFC based on a differential expression test performed in Seurat with the design. Then the average expression for each gene in the marker set within a given phase was calculated. The top 5 most highly expressed genes from each phase that had passed the differential expression filtering step were chosen as candidates for further analysis. The expression of this small set of genes was examined manually to ensure there was no cell-type-specific bias. Finally, the most strongly expressed candidates from this set were chosen for probe design. Genes from these sets that had either unknown function or were not previously characterized as being cell cycle regulated were prioritized. Probe design was performed by Molecular Instruments. *In situ* hybridization was performed as described previously^89^ with the minor modification of eliminating the proteinase K digestion to preserve the integrity of the Arabidopsis root for imaging.

### Quantification and Statistical Analysis

For scRNA-seq statistical analysis, differential expression tests to identify markers were performed using Seurat in R and the results of that statistical test are reported in Table S1. For imaging and regeneration data, statistical tests are reported throughout the manuscript and are available in figure legends. All statistical tests were performed in R. Statistical tests for data comprised of count variables were performed using the Wilcoxon test implemented in the rstatix package. Where noted, count data was tested using the Chi-square test with the stats package. Statistical tests of data comprised of continuous variables was performed the rstatix using the pairwise t-test function. The log rank test was used to determine the significance of the G1 survivorship analysis^87,88^. Loess regressions are shown throughout the manuscript with 95% confidence intervals calculated by the ggplot2 smooth function. Wherever n is less than 30, results are plotted as a combined box and jitter plot so that the n number is visible in the summary plot. Where n is greater than 30, the n value is annotated onto the summary plot.

Where fluorescence results are quantified, they are represented as the corrected total cellular fluorescence (CTCF) where the area of the relevant region of interest (ROI) was multiplied by the average fluorescence intensity of the background signal of the image. This value was then subtracted from the integrated density value of the ROI. Each of these values was obtained in FIJI using the measure function. ROIs were either determined manually based on the expression domain of a reporter gene, or were determined with automatic segmentation for all visible nuclei using either TrackMate or Imaris. In the case of *in situ* imaging experiments, ROIs were determined by manually segmenting cells based on the DAPI counterstain channel. Images were then thresholded to remove background, and the corrected total cell fluorescence (CTCF) was calculated within each ROI as described above. Cut offs to separate cells with signal from background were determined using change point analysis^83^. Permutation and bootstrap tests to determine the p-value and the confidence interval of the anti-correlation were performed in R.

Gene ontology enrichments were determined using the gene list analysis portal in Thalemine.

## Supplemental Table Titles and Legends

**<u>Table S1.</u> Summary of all differentially regulated genes identified in this study; Related to Figure 1 and S1-S5.** KmeansClust refers the cluster identified in S4. Sequencing method indicates which method the gene was detected in. sc_log2foldchange refers the log2 fold change in the scRNA-seq phase marker identification analysis. Similarly, sc_pval_adj, sc_phase, and sc_diffpct refer to the adjust p-value, enriched library, and difference in percent cells expressing in the same analysis. Marker indicates which phase a gene was identified to be a marker of for the top 50 markers. Permissive marker is the same, but includes the top 200 markers.

**<u>Table S2.</u> Gold standard markers from prior transcriptional studies; Related to Figure 1**.

**<u>Table S3.</u> Gene Set Enrichment analysis results for the top 50 and top 200 marker sets as well as the G1 bulk RNAseq clusters; Related to Figures 1 and S4.**

**<u>Table S4.</u> Differential expression analysis of G1 subpopulations; Related to Figure 2**. In the column titled “cell_group”, left refers genes with upregulated expression in the leftmost branch of cells shown in Figure 2C and upper refers to genes with upregulated expression in the uppermost branch of cells shown in Figure 2C.

**<u>Table S5.</u> G1 duration summary; Related to Figure 3**.

**<u>Table S6.</u> Cell type specific phase marker matrix. Related to Figure 2**. Each column represents a cell type plus a cell cycle phase category. Each row represents a gene. A value of 1 indicates a given gene (row) is a marker for a phase in a particular cell type (column). A value of 0 indicates a gene is not a marker of a cell cycle plus phase type.

## Supplemental Movie Titles and Legends

**<u>Movie S1.</u> Time lapse movie showing G2/M duration during homeostatic growth; Related to Figure 2**. A median section from a time lapse is shown for the *pCYCB1;1:NCYCB1;1-YFP* component of the PlaCCI reporter to show the behavior of dividing cells. Time stamp is shown in days:hr:min. CYCB1;1-YFP is shown in yellow.

**<u>Movie S2.</u> Time lapse movie showing two replicates of PlaCCI crossed to WIP4 during homeostatic growth; Related to Figure 3**. Time stamp is shown in days:hr:min. *pCYCB1;1:NCYCB1;1-YFP* is shown in yellow, *pCDT1a:CDT1a-CFP* is shown in cyan and *pHTR13:HTR13-mCherry* is shown in red. The *pCDT1a:CDT1a-CFP* channel is also shown in a separate panel. Panels A and B show two replicates of time lapses taken under control conditions of roots undergoing homeostatic growth.

**<u>Movie S3.</u> Time lapse movie showing two replicates of PlaCCI crossed to *pWIP4:GFP*, *PET111:YFP*, or *pWOX5:YFP* during regeneration; Related to Figure 3**. Time stamp is shown in days:hr:min and represents time post ablation. *pCYCB1;1:NCYCB1;1-YFP* is shown in yellow, *pCDT1a:CDT1a-CFP* is shown in cyan and *pHTR13:HTR13-mCherry* is shown in red. The *pCDT1a:CDT1a-CFP* channel is also shown in a separate panel. PlaCCI is shown in each panel and additional reporters are shown as follows: (A) *PET111:YFP*, (B) *pWOX5:YFP*, (C) *PET111:YFP*, (D) *pWIP4:GFP*. In panels A, B, and C the following reporters are shown in yellow in addition to CYCB1;1: in the top and bottom panels on the left, PET111 is shown in yellow. In the top right panel, WOX5 is shown in yellow. Panel D shows WIP4*pWIP4:GFP* expression in grayscale as a separate panel.

**<u>Movie S4.</u> Time lapse showing GSH burst following an ablation; Related to Figure 4**. Time stamp is shown in days:hr:min where 00:00:00 marks the first frame of the time lapse. *pCYCB1;1:NCYCB1;1-YFP* is shown in yellow, pCDT1a:CDT1a-CFP is shown in cyan and pHTR13:HTR13-mCherry is shown in red, and Blue CMAC is shown in grey. The Blue CMAC channel is also shown in a separate panel.

